# Integrative inference of transcriptional networks in Arabidopsis yields novel ROS signalling regulators

**DOI:** 10.1101/2020.08.11.245902

**Authors:** Inge De Clercq, Jan Van de Velde, Xiaopeng Luo, Li Liu, Veronique Storme, Michiel Van Bel, Robin Pottie, Dries Vaneechoutte, Frank Van Breusegem, Klaas Vandepoele

## Abstract

Gene regulation is a dynamic process in which transcription factors (TFs) play an important role to control spatiotemporal gene expression. While gene regulatory networks describe the interactions between TFs and their target genes, our global knowledge about the complexity of TF control for different genes and biological processes is incomplete. To enhance our global understanding of regulatory interactions in *Arabidopsis thaliana*, different regulatory input networks capturing complementary information about DNA motifs, open chromatin, TF binding and expression-based regulatory interactions, were combined using a supervised learning approach, resulting in an integrated gene regulatory network (iGRN) covering 1,491 TFs and 31,393 target genes (1.7 million interactions). This iGRN outperforms the different input networks to predict known regulatory interactions and has a similar performance to recover functional interactions compared to state-of-the-art experimental methods like yeast one-hybrid and ChIP-seq. The iGRN correctly inferred known functions for 681 TFs and predicted new gene functions for hundreds of unknown TFs. For regulators predicted to be involved in reactive oxygen species stress regulation, we confirmed in total 75% of TFs with a function in ROS and/or physiological stress responses. This includes 13 novel ROS regulators, previously not connected to any ROS or stress function, that were experimentally validated in our ROS-specific phenotypic assays of loss- or gain-of-function lines. In conclusion, the presented iGRN offers a high-quality starting point to enhance our understanding of gene regulation in plants by integrating different experimental data types at the network level.

## INTRODUCTION

Accurate spatiotemporal gene expression is of key importance for plant development and molecular responses to environmental cues. Key factors controlling this process are transcription factors (TFs) which regulate their target genes by recognizing and binding short *cis*-regulatory DNA sequences called TF binding sites (TFBSs or DNA motifs) ^1^. The full set of regulatory interactions between TFs and their target gene(s) forms a gene regulatory network (GRN) and depending on the underlying source of data, GRNs are composed of physical and/or regulatory protein-DNA interactions. Efficiently identifying regulatory interactions is pivotal to understand how a gene’s transcriptional activity is controlled through the interplay of one or multiple regulators. Driven by different modes of gene duplication, specific TF families have been expanded within the green plant lineage, with the model species *Arabidopsis thaliana* containing around 1700 TFs ^2^. Although specific regulatory networks and the associated TFs have been studied and are often conserved across species ^3^, still for hundreds of *Arabidopsis TFs* experimental information is missing ^4^.

Several studies have used high-throughput assays to systematically identify GRNs under specific conditions or in specific developmental and/or anatomical contexts in *Arabidopsis*. Examples include yeast one-hybrid (Y1H) screens to map process-specific GRNs as well as TF chromatin immunoprecipitation (ChIP) experiments describing regulatory interactions between TFs and target genes involved in flower development, light and clock signaling, hormone response, and (a)biotic stress response ^5^. ChIP data revealed the high complexity of plant gene regulation and suggested that a single gene can potentially be regulated by up to 75 different TFs ^6,7^. Open chromatin profiling, either using DNase-Seq or Assay for Transposase-Accessible Chromatin using sequencing (ATAC-Seq), allows studying the organization and dynamics of accessible chromatin regions, and provides a template to infer GRNs in different organs, in response to specific treatments or in different cell types ^8-12^. Whereas all these methods aid the construction of GRNs by reporting physical protein-DNA interactions, a major challenge remains the identification of functional interactions, referring to the binding of a specific TF to a cis-regulatory sequence leading to a regulatory effect on a target genes’ expression.

In this paper we present a network-based approach based on supervised learning for large-scale functional data integration to enhance our understanding of TF gene regulation in *Arabidopsis*, hence referred to as an integrative GRN (iGRN). Supervised learning approaches make efficient use of the available prior knowledge to train a machine learning algorithm, called a classifier, for handling new data. Compared to unsupervised learning approaches, supervised learning is considered more powerful when a large amount of existing knowledge is available to guide the classification task, which here is the identification of functional TF-target interactions^13^. Construction and validation of the iGRN, leveraging a diverse set of complementary experimental datasets, revealed a strong enrichment for functional interactions and a high predictive power to correctly infer functions for both functionally characterized and uncharacterized TFs involved in a variety of biological processes. The latter is demonstrated in a case study by the experimental validation of novel iGRN-based function predictions for TFs involved in plant responses to oxidative stresses.

## RESULTS

### An integrated gene regulatory network for *Arabidopsis thaliana*

To generate an integrated gene regulatory network (*Arabidopsis* iGRN), different types of complementary regulatory datasets were first processed, resulting in different input networks (Table 1). In total seven input networks, encompassing information about DNA motifs, open chromatin, TF binding and expression-based regulatory interactions, were generated by applying different network inference and data analysis approaches (see Methods for underlying datasets; Figure 1a). Subsequently, through the application of a supervised learning approach, a classifier was built to identify those interactions from the input networks that most likely represent functional interactions.

**Table 1.** Overview of different networks used to construct the iGRN.

| Type | Network name | #interactions | #TFs | Median #interactions (average) |
| --- | --- | --- | --- | --- |
| Input network | PWM | 17,053,785 | 874 | 21,797 (19,512) |
|  | Conserved PWM | 340,893 | 874 | 247 (390) |
|  | PWM in DHS | 5,231,258 | 874 | 6,195 (5,985) |
|  | COE + PWM | 324,864 | 703 | 290 (462) |
|  | TF ChIP | 46,619 | 27 | 1,026 (1,727) |
|  | COE TF-target | 1,547,002 | 1547 | 1,000 (1,000) |
|  | GENIE3 | 2,588,312 | 1261 | 1,985 (2,053) |
| Gold standard | Gold standard | 5,732 | 522 | 2 (11) |
| Evaluation | Test dataset | 1,146 | 257 | 1 (4) |
| Output network | iGRN | 1,709,871 | 1491 | 802 (1,147) |

**Figure 1.**
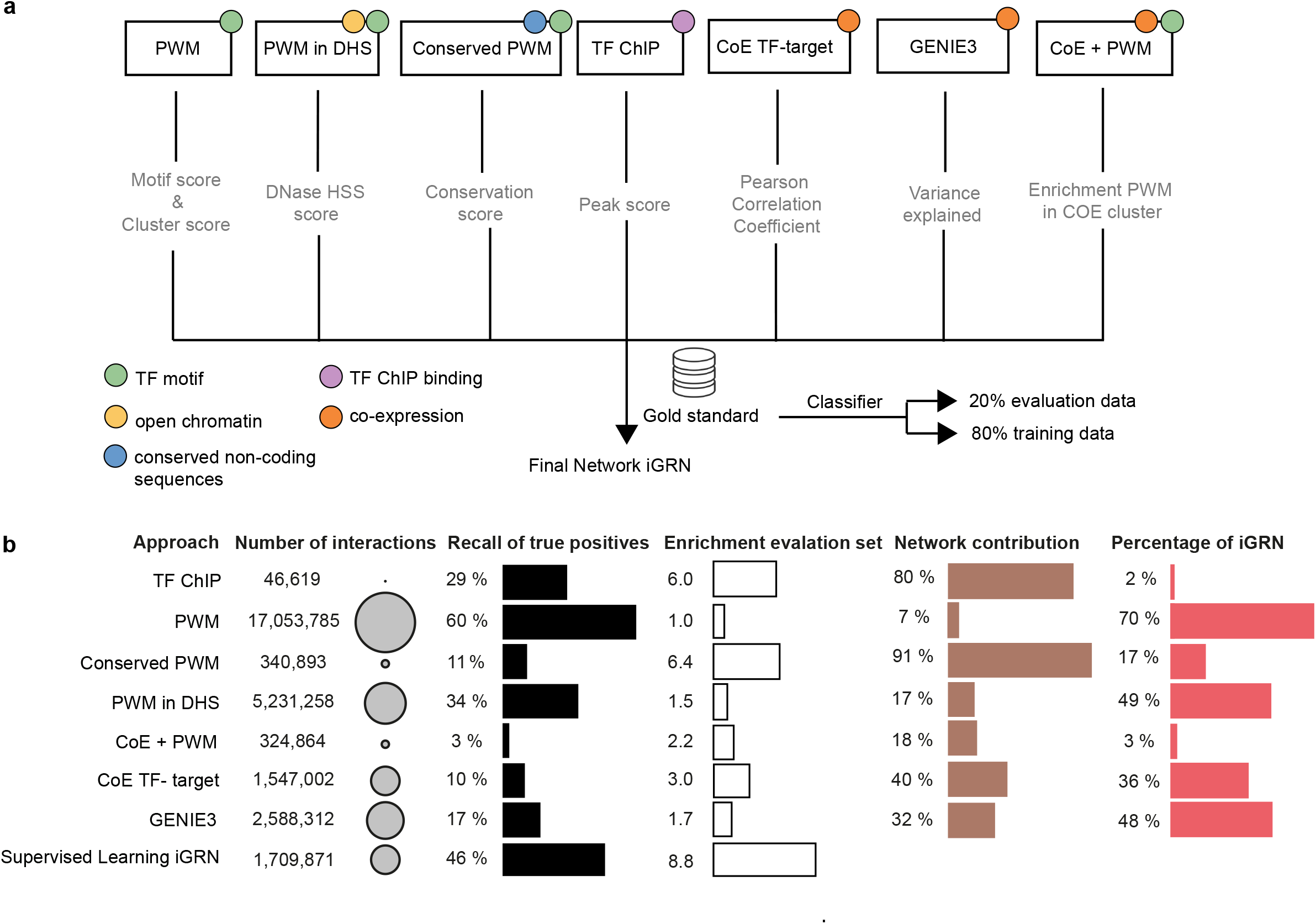
Construction and evaluation of the integrated gene regulatory network (iGRN). a. Overview of the different input networks used to construct the supervised iGRN. Whereas colored circles indicate the type of data that is used to construct the input network, the information below each network depicts how edges in that network are weighted. b. Overview of the sizes of the different networks together with the recall and enrichment for edges in the evaluation test dataset (black and white bars, respectively). The brown bars report the fraction of each input network that is present in the iGRN while the red bars report which fraction of the supervised iGRN edges is supported by this input network.

The first network is based on the presence of a TF motif, modelled using Position Weight Matrices (PWMs), in the promoter of a target gene. A total of 1,260 high quality PWMs, covering 874 TFs, were mapped onto the *Arabidopsis* genome resulting in a TF - PWM target gene network (called “PWM”). Using the genomic locations of publicly available DNaseI datasets the PWM network was filtered resulting in a second open chromatin regulatory input network (called “PWM in DHS”). DNaseI hypersensitive sites (DHS) are considered to be a proxy for open chromatin and potential protein binding to the DNA ^14^. As a third network, the set of PWMs was also evaluated for evolutionary conservation across 13 dicotyledonous genomes using a comparative motif mapping algorithm ^15^, which resulted in an evolutionarily conserved regulatory network (“Conserved PWM”). To integrate experimental data, a fourth “TF ChIP” network was created by combining experimentally determined TF binding occupancy profiles from 27 TFs ^7^. Next to a physical TF binding network, two co-expression-based regulatory networks were also generated. Co-expression approaches measure similarity at the output level of transcriptional regulation and as such can provide a complementary view compared to promoter-centered regulatory networks. First, the Pearson correlation between a TF and a target gene was used to build co-expression clusters around each TF, resulting in a fifth “CoE TF-target” input network. Secondly, GENIE3, a reverse engineering tool for reconstructing regulatory networks based on expression data, was used to build a sixth input network ^16^. GENIE3 was among the best performing tools in reconstructing regulatory interactions in a large-scale benchmark study ^17^. The last network was generated using a hybrid approach of co-expression and PWM based methods (“COE+PWM”). Initially a co-expression cluster was constructed for each gene and PWM enrichment was subsequently performed. Only genes with significantly enriched PWMs in their co-expression cluster were retained, resulting in a seventh network where TF binding site information is combined with co-regulatory information (see Methods).

As the different input networks capture different aspects of gene regulation, we next applied supervised learning to construct, starting from these input networks, an integrative regulatory network reporting functional interactions. A score for each possible TF-target interaction was devised as a function of the feature-specific weights provided by the seven input networks. For building a classifier, we compiled a gold standard dataset comprising interactions based on all validated interactions in AtRegNet ^18^, an exhaustive literature-based network reporting interactions in secondary cell wall development ^19^, and a large-scale data mining effort on 974 peer reviewed articles ^20^. This gold standard entails interactions for 47 TF families involved in different biological processes, with on average 11 interactions per TF (Table 1 and Supplemental Table S1). Eighty percent of the interactions present in this gold standard were retained as training dataset for the machine learning classification algorithms while the remaining twenty percent was used as an evaluation set. Initially different classifiers were learned and their performance to identify functional interactions in the evaluation set was assessed (Supplemental Table S2). The classifier based on gradient boosting model using decision trees showed the best performance, with an area under the curve (AUC) value of 0.77, and Precision and Recall of 71% and 46% on the evaluation set, respectively (Supplemental Table S2). Applying this classifier to the complete set of interactions present in the seven input networks resulted in an integrative network of 1,709,871 interactions for 1,491 TFs, containing 31,393 target genes (29,137 protein-coding and 2,256 non-coding genes; see Data file 1).

In order to obtain an unbiased performance estimate of the input networks compared to the iGRN, all input networks were compared to the evaluation dataset of known interactions (20% of the gold standard) and benchmarked measuring the recovery of true edges (referred to as the recall, Figure 1b). Furthermore, we also determined the enrichment of a network towards the evaluation dataset, which functions as a proxy for specificity ^21^. Focusing on the different input networks, Figure 1b shows that the network based on evolutionary conserved TF binding sites (“Conserved PWM”) has the highest enrichment (6.4 fold, Supplemental Table 3). Evolutionary conservation is known to be a strong predictor of functional regulatory elements explaining this high enrichment ^22,23^. The TF ChIP-seq network also performs well with both a high recovery (29%) and enrichment (6.0 fold). This overlap can partially be explained by the fact that a subset of the validated AtRegNet interactions in the gold standard are based on ChIP studies ^7^. Other TF motif mapping strategies recover larger fractions of true edges but suffer from low specificity, resulting in low enrichment values. The simple mapping of TF motifs (“PWM”) is known to result in many false positives because TF binding sites are often short and typically contain some level of degeneracy ^11,24^. Both co-expression based networks do not consider TF binding site information and are therefore able to also identify regulatory interactions for TFs lacking DNA binding motif information. Both these networks perform relatively well: correlation-based co-expression between TF and target genes, implemented in the “CoE TF-target” network, has the third highest enrichment towards the evaluation dataset (10% recovery and 3.0 fold enrichment), while GENIE3 displays a good balance between recovery of true edges and specificity (Supplemental Table S3). Focusing on the integrative network (iGRN) shows it has the strongest enrichment (8.8 fold) as well as a high recovery for true regulatory relationships (46% recovery of interactions in the evaluation set), clearly outperforming the different computational and experimental approaches used to construct the input networks.

Apart from determining the power of the iGRN network to recover known functional interactions, we also measured the contribution of each of the seven input networks to the iGRN. Ideally, a good integration approach should favor interactions from the input networks with high enrichment values towards the evaluation set, as these would boost the recovery of correct interactions. As reported in Figure 1b (brown and red bar charts), the evolutionary conserved motifs (“Conserved PWM”) and the “TF ChIP” network contributed the largest part of their edges to the integrative network (91% and 80%, respectively), while networks showing low specificity, such as “PWM”, only contributed a small fraction of interactions (7%). This result corroborates that the applied supervised approach successfully identified functional interactions coming from high-quality input networks. Next, we measured which iGRN interactions are supported by the different input networks (Figure 1b, red bar plot). Overall, the integrative network has 70% of the edges supported by a PWM mapping, meaning that most regulatory interactions are supported by the presence of TF binding site(s). Furthermore, 36-48% of the interactions are supported by at least one of the two co-expression networks, indicating for nearly half of all interactions a regulatory effect at the transcript level. Overall, 92% of the iGRN interactions are supported by more than one input network (Figure S1), confirming both the commonality and complementarity of the input networks.

### iGRN interactions are complementary to TF regulatory events identified from experimental methods

To assess whether the iGRN identifies functional TF-target gene interactions that are supported by experimental data, detailed comparisons of the iGRN interactions were made against three different datasets, capturing a variety of TFs and biological processes, which were not used during the iGRN construction (see Methods). These datasets cover i) a set of 2,759 Y1H interactions reflecting direct *in vitro* interactions ii) 28 TF ChIP-seq experiments collected from different studies (148,949 interactions, see also Methods) reflecting primarily direct *in vivo* binding and iii) a set of 40 differentially expressed (DE) gene sets upon genetic TF perturbation (51,178 interactions, see Methods). The significance of the observed overlaps was examined through comparison to a background distribution of expected overlaps generated by randomizing the iGRN (see Methods). First, we detected a weak but significant overlap between edges in the integrative network and regulatory edges determined by Y1H (Figure 2a). An overlap of 336 edges (12%) was observed reflecting a 1.6 higher overlap than expected by chance (Figure 2b). TF ChIP-seq interactions displayed an overlap of 15,487 interactions (10%) with a fold enrichment of 2.4. Only considering TFs that occur in both the iGRN and this ChIP-Seq dataset, for 25/28 regulators there was a significant overlap between the iGRN and ChIP defined target genes (Supplemental Table S4). Overall, we found that TF ChIP-seq data confirmed 39% of predicted interactions in the iGRN, demonstrating that a large fraction of iGRN interactions are supported by *in vivo* TF binding. This fraction is comparable with the concordance between TF ChIP-based regulatory interactions and altered target gene transcript levels after genetic TF perturbation, an approach that is frequently used to identify bound target genes that are also regulated ^25^. Next to TF-DNA binding assays, we also investigated the recovery of directly and indirectly regulated target genes based on differential target gene expression. The iGRN recovered 9,672 differentially expressed interactions (19%) resulting in a 3.6 fold enrichment (Figure 2a and 2b). Taken together, both the comparison with binding and regulatory datasets demonstrate that the iGRN represents high-quality interactions supported by experimental data.

**Figure 2.**
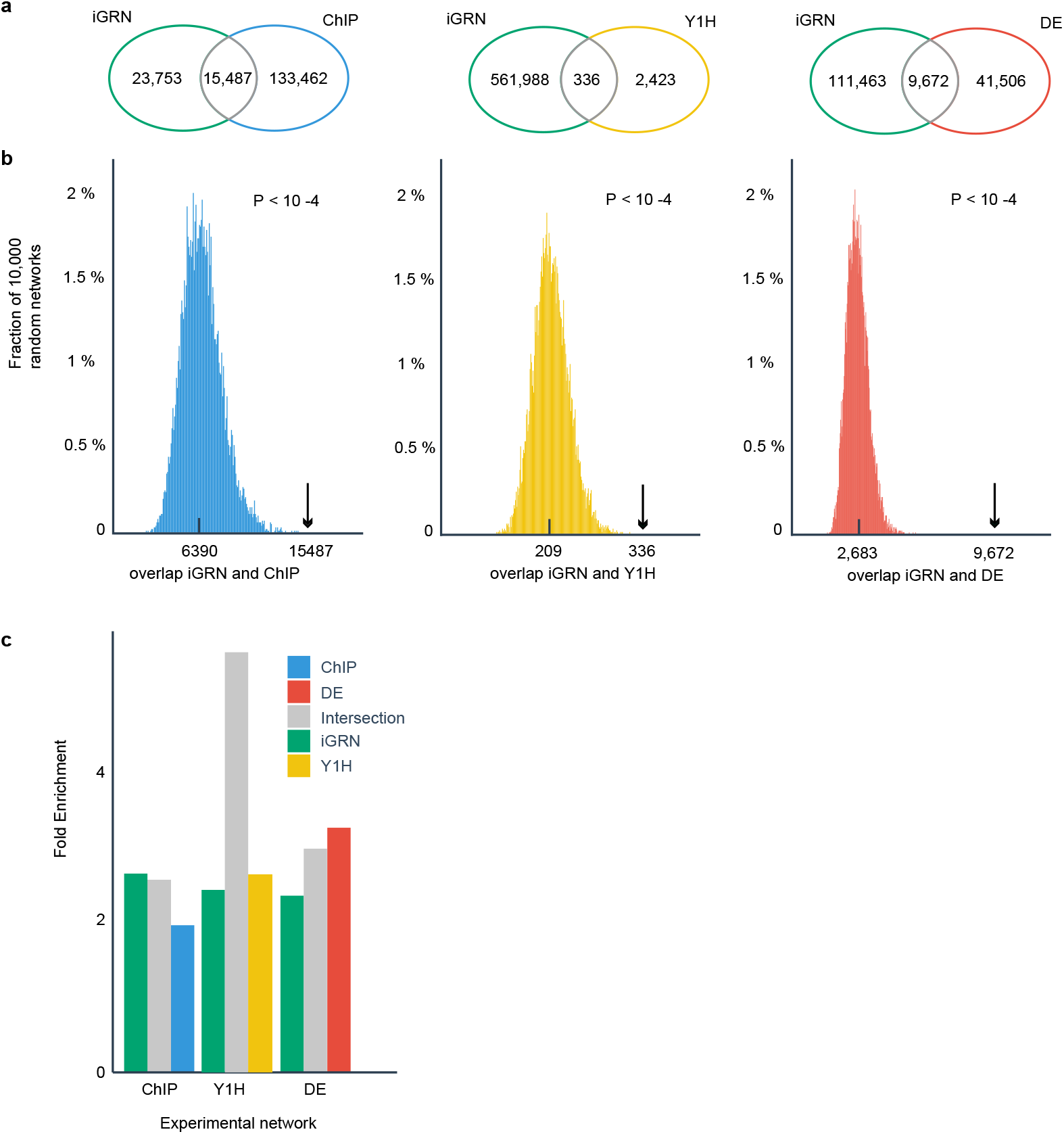
Overlap between iGRN interactions and different experimental data sources describing gene regulatory and functional information. a. Venn diagrams showing the number of shared interactions between the iGRN and TF ChIP, Y1H and differentially expressed (DE) genes after TF perturbation (depicted in blue, yellow and red, respectively). b. Histograms showing the expected overlap based on 10,000 randomized networks, together with the observed overlap indicated by the black arrow. The p-values for observing the measured overlap by chance are also reported (permutation based statistics, one-sided). c. Fold enrichment for a functional metric describing how well target genes of a specific TF are involved in similar biological processes (average values for all enriched GO terms and all TFs present in the different data series are reported). Whereas the grey bars indicate the functional enrichment for shared interactions, blue, red, yellow and green depict fold enrichment for interactions unique to the ChIP, DE, Y1H and iGRN, respectively.

To evaluate the relevance of the regulatory interactions uniquely predicted by the iGRN and hence not overlapping with the three experimental datasets (Y1H, TF ChIP-Seq, and DE), a functional metric was used to measure whether a TF’s target genes participate in similar biological processes. This metric starts from the assumption that a set of bona fide target genes will have a higher enrichment for functional annotations than randomized networks ^21^. For each TF, the enriched Gene Ontology (GO) Biological Process terms based on its target genes were determined, summarized in a distribution and compared to the background distribution based on randomized networks. In Figure 2c, the iGRN-predicted interactions were first divided in different sets based on their overlap with the TF ChIP, Y1H and DE experimental datasets (as in Figure 2a), respectively. Next, for each subset the fold enrichments for functional GO Biological Process terms were assessed. Whereas the grey series in Figure 2c show the regulatory interactions shared between the iGRN and the experimental dataset, the green series shows the interactions unique to the iGRN. Likewise, the blue, yellow and red series show the regulatory interactions present in the ChIP, Y1H and DE dataset not overlapping with the iGRN.

Focusing on the overlap with TF ChIP dataset revealed that the functional enrichment was similar for the interactions unique to the iGRN compared to those shared between the iGRN and TF ChIP dataset (2.63 and 2.55, respectively). However, the regulatory interactions unique to the TF ChIP datasets showed lower fold enrichments (1.94), indicating that these show less functional coherence. For the Y1H comparison, the interactions uniquely predicted by the iGRN had a similar fold enrichment as the interactions only reported by the Y1H assays (2.41 and 2.61). The interactions that are reported by both approaches had a considerable higher enrichment compared to the other networks (5.55; though note the small set size, n=336). The comparison with the DE dataset showed a similar result with an increase in enrichment fold from uniquely predicted edges in the iGRN to the intersection and uniquely predicted edges by the DE gene sets (2.33, 2.96 and 3.23, respectively). Taken together, these results indicate that the predicted interactions, including those unique to the iGRN, convey functional gene regulatory events.

### Network architecture reveals different levels of gene regulatory complexity affecting plant phenotypes

Starting from the complete iGRN comprising 1.7 million interactions, we investigated the structure and complexity of the network and determined if network architecture can be linked to specific biological functions^26^. On average, each gene in the network has 54 incoming TFs (median 32), while each TF is linked to 1,147 target genes (median 802; Supplemental Figure S2). There was a large range in connectivity with half of the genes being regulated by 8-79 TFs (interquartile range), and a small number of TFs (107/1491, or 7%) regulating more than 3,000 target genes. Focusing on the top 1% genes with the largest number of incoming TFs revealed a significant bias towards genes with transcription factor activity, corroborating previous findings reporting that regulators themselves are complexly regulated by other regulators ^7,27^. In contrast, 2172 genes having only one incoming TF showed enrichment for RNA binding and translation, suggesting that these functions are less complexly regulated by TFs.

Next, we examined if network structure can offer new insights in the role of individual regulators. Concretely, we investigated whether the degree (number of target genes) and betweenness centrality (a measure for the importance based on number of shortest paths in the network graph) of TFs in the iGRN can be predictive for potential master regulators showing prominent phenotypes upon genetic perturbation. Whereas degree refers to the number of interactions a node, or gene, has in the network, betweenness centrality measures how important a node is to the shortest paths through the network. Nodes with a high centrality are assumed to be more important in the network as more information is passing through these nodes. After processing large-scale phenotyping data from RARGE II, an integrated phenotype database of Arabidopsis mutants scoring traits using a controlled vocabulary for 14,540 genes, phenotypes were recorded for 2,224 genes ^28^. When splitting all TFs present in the iGRN and represented in the RARGE II dataset in four quartiles (Q1-Q4) based on their degree values, we observed that 17.4% of the high-degree (Q3+Q4) and 15.1% of the low-degree (Q1+Q2) TFs had a reported phenotype (p-value 0.43; two-tailed Mann-Whitney U test). When comparing the quartiles of TFs based on betweenness centrality, a significant difference was found when scoring regulators with a reported phenotype (19.6% for Q3+Q4 and 12.4% for Q1+Q2, respectively; p-value 0.0088; two-tailed Mann-Whitney U test; Figure 3a). This pattern is confirmed when comparing the betweenness centrality for TFs with and without phenotypes, as regulators showing RARGE II phenotypes have more of the shortest network paths going through their nodes (log2 median value of 17.50 and 16.46 for TFs with and without reported phenotypes, respectively; p-value 1e-05; two-tailed Mann-Whitney U test; Figure 3b). iGRN TFs within the top 10% genes with highest centrality for which mutants have reported phenotypes (19/91) are significantly biased towards post-embryonic development and include ATGTL1, PIF4, SEP3, IDD4 and IDD8, ANAC056, DOF5.6, LFY3 and TCP4. After assigning regulators to three levels based on their network hierarchy ^29^, we observed that top-level TFs high in the hierarchy (n=313) have higher degree and centrality values compared to bottom-level TFs (n=704, median degree for top and bottom TFs is 1923 and 529, respectively, while log2 median betweenness centrality for top and bottom TFs is 18.87 and 16.10, respectively). Top-level TFs were not significantly enriched for specific TF families. Interestingly, both for TFs with and without phenotypes, a similar number of TFs in their target genes was found (9.45% and 9.44%, respectively), indicating that TFs showing a phenotype do not regulate more TFs and thus are not higher in the TF hierarchy.

**Figure 3.**
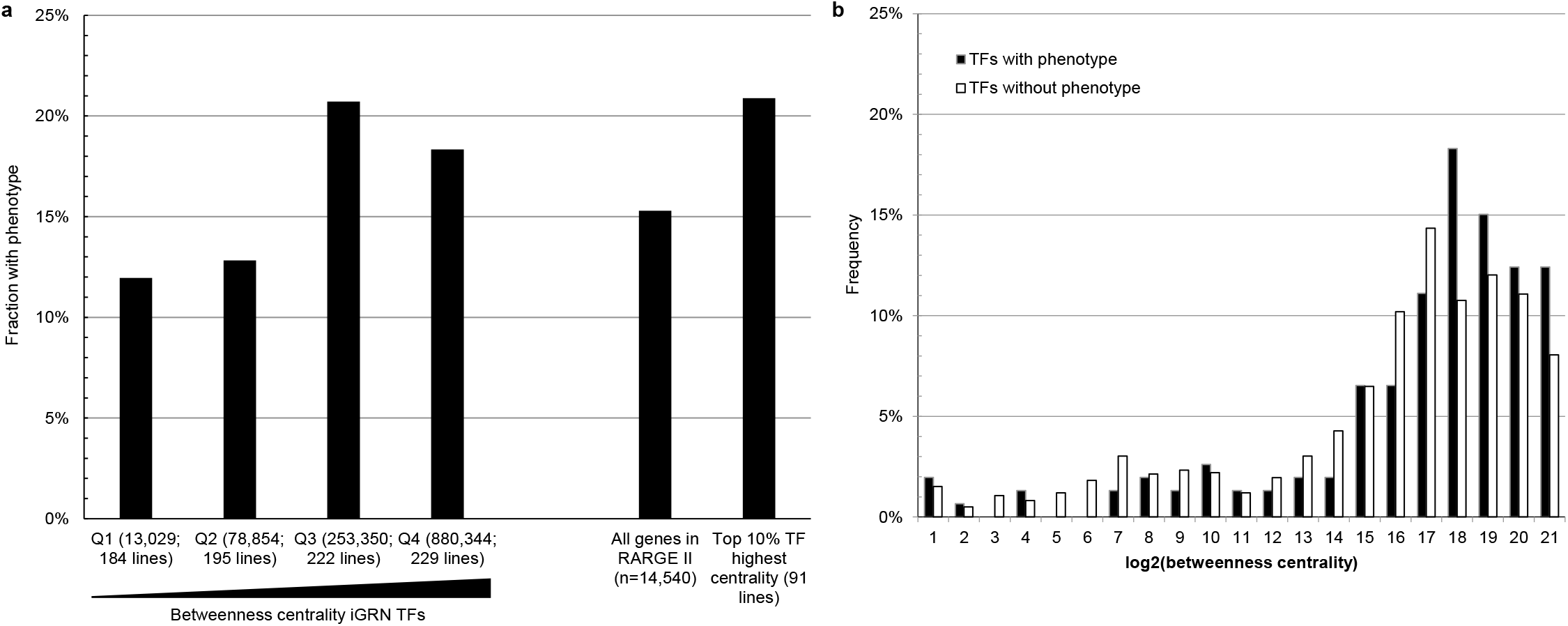
Phenotypes reported after TF perturbation for TFs with different network centrality properties. a. RARGE-II phenotypes reported for iGRN TFs with different betweenness centrality. Q1-Q4 refer to the four quartiles of all iGRN TFs (sorted based on increasing betweenness centrality) and values in parenthesis refer to the median betweenness centrality value together with the number of lines available in the RARGE II dataset for that quartile. b. Betweenness centrality distribution for TFs with and without phenotypes reported in RARGE-II (n= 153 and 1589 TFs, respectively).

As transcriptional regulation is mediated through interactions between TFs, co-activators or co-repressors, and the general transcriptional machinery, we determined whether the iGRN has predictive power to identify cooperative TF regulation. Therefore, we measured if regulators sharing a large number of target genes are more likely to interact with each other, based on experimentally determined protein-protein interactions (PPIs). We found that TF pairs sharing many target genes (Jaccard index >0.3) have a 15-fold higher occurrence of PPIs compared to TF pairs having less shared target genes (Jaccard index <0.3), which is highly significant (2.71% versus 0.31%; p-value <2.2e-16, hypergeometric distribution). Whereas most of these PPIs cover TFs from the same gene family, probably due to the high similarity of their corresponding binding sites yielding similar sets of target genes, their potential to act as heterodimers and/or potential functional redundancy, also cases of interacting TF pairs from different gene families were found. Examples of the latter case include HY5-PIF3/PIF4 involved in far-red signaling, MYB21-RGL1/RGL2 controlling hormone-mediated cell proliferation and differentiation, and BES1-BIM1/BIM2 acting synergistically to control brassinosteroid signaling (Supplemental Figure S3). The highly correlated expression profiles of these TFs sharing many target genes further suggest they exert their regulatory roles in a cooperative manner.

### Network-based identification of gene functions for Arabidopsis TFs

Based on the validation results indicating the functional relevance of regulatory edges in the iGRN, we also investigated if individual TF functions can be inferred. Therefore, we performed GO enrichment analysis for all target genes of each TF, only considering experimental Biological Process annotations and excluding very general GO terms (see Methods). Next, we determined if the enriched Biological Process terms were already known for that specific TF or whether a novel functionality was predicted (Data file 2. TF_function_all). In total we could infer functions for 1,301 TFs and for 54% (681) of the 1,259 TFs with experimental Biological Process annotations we could correctly infer one or more of the known functions. For 268 TFs lacking experimental annotations, novel iGRN-based functions were predicted. Reciprocally, per Biological Process term, on average 21% of the known regulators were recovered, while for several distinct processes the recovery was 50% or higher. Examples of processes with high recovery rates include response to wounding (14/20, 70%), plant-type secondary cell wall biogenesis (18/28, 64%), response to hormone (174/343, 51%), chloroplast organization (6/12, 50%) and DNA replication (3/6, 50%).

The overall recovery of known TF functions based on the iGRN was lower compared to the probabilistic functional gene network AraNet v2 ^30^, which recovered known functions for 739 TFs (28% average recovery of known regulators per GO BP term). However, both approaches are highly complementary, as 207 and 265 TFs had known functions only recovered using iGRN and AraNet, respectively (Supplemental Figure S4). Examples of known TF functions uniquely recovered through the iGRN cover EIN3 (ethylene-activated signaling pathway), APETALA2 (flower development), LFY (flower development, gibberellic acid-mediated signaling pathway), FUS3 (somatic embryogenesis), TINY (multidimensional cell growth), and AMS, ATWRKY34 and CRF12 (pollen development). Focusing on specific GO terms, biological processes with a higher recovery of known regulators in the iGRN compared to AraNet cover response to biotic stimulus (47% versus 35%), chloroplast organization (50% versus 0%), response to abiotic stimulus (47% versus 26%), pollination (33% versus 0%), and root hair cell differentiation (35% versus 6%). For the latter process we correctly identified the regulators ZPF5, RSL2 and RSL4, RHD6, LRL2 and LRL3, as well as MYB36. MYB36 regulates the developmental transition in the root endodermis, where the differentiated endodermis forms a protective waxy barrier called the Casparian strip. In our network, MYB36 is predicted to regulate genes involved in cell wall organization such as CASP1-5, PER64 and ESB1, which are known to be involved in Casparian strip formation and have been shown to be differentially expressed in the *myb36* mutant ^31^. Furthermore, the iGRN also predicts MYB36 to be involved in root hair elongation, which agrees with the longer root hairs observed in *myb36* mutants ^31^.

### Identification of novel TFs in reactive oxygen species (ROS) stress regulation

Apart from recovering known functions of TFs involved in development and stress responses, the predictive power of the iGRN was used to identify novel TF functions. As a test case, we extracted TFs from the iGRN with predicted oxidative stress signaling and/or responsiveness functions. As an increase in ROS levels is a common theme during abiotic and biotic stresses, oxidative stress responsiveness was used to validate novel regulators of plant stress responses. One hundred twenty-four ROS-related TFs were extracted from the iGRN by identifying those TFs that entailed an enrichment for core ROS responsive transcripts (ROS marker genes) in their target genes ^32^. Among these TFs, 17 had an already reported function in plant responses to ROS or ROS-causing agents (hydrogen peroxide [H_2_O_2_], methyl viologen [MV] and 3-amino triazole [3-AT]), here referred to as ‘KNOWN_ROS’ TFs (Table 2, Supplemental Table S5). Fifty-eight ROS TFs were previously reported to function in plant responses to environmental stresses, such as salinity, drought, heavy metals, cold, hypoxia, high light and infection by bacterial and fungal pathogens (and that we refer to as ‘KNOWN_STRESS’ TFs, Supplemental Table S5). The remaining 49 identified ROS TFs had previously not been linked to any role in ROS/environmental stress (referred to as ‘NOVEL’ TFs in Table 2), at the time the iGRN was constructed (see Methods). However, five of the 49 novel TFs were validated for a role in environmental stress responses by studies that were published after the iGRN construction (Table 2 and Supplemental Table S5): ANAC102 and BME3 (high light ^33-35^), NAC103 (genotoxic ^36^), ZAT18 (drought ^37^) and ERF019 (drought, bacterial and fungal pathogens ^38,39^).

**Table 2.** Overview phenotype information for known and validated novel ROS regulators.

| **RANK**<br>**iGRN** | **Symbol** | **q-val** | **Status (1)** | **Known**<br>**oxidative**<br>**stress**<br>**phenotype**<br>**from**<br>**literature** | **Reference** | **Known other**<br>**(a/biotic) stress**<br>**phenotype from**<br>**literature** | **Reference** | **Validated**<br>**oxidative stress**<br>**phenotype from**<br>**this study** |
| --- | --- | --- | --- | --- | --- | --- | --- | --- |
| **iGRN1** | ANAC102 | 1.84E-35 | **NOVEL** |  |  | high light | <sup>33</sup> | **MV** |
| **iGRN3** | ZAT11 | 2.81E-25 | KNOWN_ROS | MV | <sup>85</sup> | heavy metal | <sup>86</sup> |  |
| **iGRN5** | ZAT12 | 8.70E-25 | KNOWN_ROS | MV | <sup>87</sup> | osmotic/salt/high<br>light/heat | <sup>88</sup> |  |
| **iGRN6** | WRKY30 | 1.42E-24 | KNOWN_ROS | MV | <sup>89</sup> | salt | <sup>89</sup> |  |
| **iGRN11** | HSFA4A | 5.09E-23 | KNOWN_ROS | MV | <sup>90</sup> | salt, anoxia | <sup>90</sup> |  |
| **iGRN12** | WRKY15 | 7.46E-23 | KNOWN_ROS | H2O2 | <sup>91</sup> | salt/osmotic | <sup>91</sup> |  |
| **iGRN16** | STZ | 4.90E-22 | KNOWN_ROS | MV | <sup>92</sup> | osmotic/salt/heat | <sup>92</sup> |  |
| **iGRN19** | SCL13 | 3.76E-21 | **NOVEL** |  |  |  |  | **MV** |
| **iGRN22** | MYB51 | 1.84E-20 | **NOVEL** |  |  |  |  | **3-AT, MV** |
| **iGRN27** | BME3 | 2.01E-18 | **NOVEL** |  |  |  |  | **MV** |
| **iGRN29** | WRKY55 | 6.69E-18 | **NOVEL** |  |  |  |  | **3-AT, MV** |
| **iGRN36** | ERF6 | 3.13E-15 | KNOWN_ROS | H2O2 | <sup>93</sup> | high light, B.<br>cinerea, osmotic | <sup>93 94 95</sup> |  |
| iGRN50 | WRKY45 | 5.93E-13 | NOVEL(*) |  |  | submergence stress | 47 | 3-AT, MV |
| iGRN52 | ANAC032 | 1.26E-12 | KNOWN_ROS | 3-AT | 96 | P. syringae, salt/high light | 96 97 |  |
| iGRN55 | AT2G33710 | 9.71E-12 | NOVEL |  |  |  |  | 3-AT, MV |
| iGRN58 | MYB122 | 1.42E-10 | NOVEL |  |  |  |  | 3-AT, MV |
| iGRN60 | WRKY28 | 1.70E-10 | KNOWN_ROS | MV, H2O2 | 98 | salt/osmotic/drought, S. sclerotiorum, A. brassicicola | 98 99 100 |  |
| iGRN62 | ANAC013 | 1.93E-10 | KNOWN_ROS | MV | 101 |  |  |  |
| iGRN65 | ABF3 | 8.06E-10 | KNOWN_ROS | MV | 102 | cold/freezing/heat/drought | 102 |  |
| iGRN66 | ERF022 | 9.55E-10 | NOVEL |  |  |  |  | MV |
| iGRN68 | BEH2 | 2.75E-09 | NOVEL |  |  |  |  | 3-AT, MV |
| iGRN69 | ANAC017 | 3.04E-09 | KNOWN_ROS | MV | 103 | drought/high light | 104 |  |
| iGRN81 | PHL1 | 2.76E-08 | NOVEL |  |  |  |  | 3-AT, MV |
| iGRN85 | JAM1 | 4.07E-08 | KNOWN_ROS | MV, H2O2 | 98 | salt/osmotic/drought | 98 |  |
| iGRN93 | MYC2 | 4.19E-07 | KNOWN_ROS | MV | 105 | salt, B. Cinerea | 106 107 |  |
| iGRN100 | AT5G57150 | 1.50E-06 | NOVEL |  |  |  |  | 3-AT, MV |
| iGRN102 | ERF115 | 1.73E-06 | NOVEL |  |  |  |  | 3-AT, MV |
| iGRN110 | JUB1 | 3.20E-06 | KNOWN_ROS | H2O2 | 108 | salt/heat, P. syringae, drought | 108 109 110 |  |
| iGRN111 | HSFA2 | 3.21E-06 | KNOWN_ROS | MV | 111 | high light, heat, cold, hypoxia | 111 112 113 114 115 |  |
| iGRN118 | CBF1 | 5.32E-06 | KNOWN_ROS | MV | 116 | salt, drought, freezing | 116 117 118 |  |

(1) Asterisk indicates NOVEL TFs that were validated for a function in (a)biotic stress (\*) responses by literature evidence after 2016.

For experimental validation, we selected the top 32 novel TFs, ranked based on enrichment for ROS marker target genes^32^, for which at least one *Arabidopsis* loss- or gain-of-function transgenic line was available (Supplemental Table S6). Plant performance under oxidative stress was assessed based on rosette growth of the mutant lines compared to the wild type when grown at low and high MV and 3-AT concentrations, respectively, using automated image analysis (see Methods), and, in addition, chlorosis symptoms were visually scored.

For 13 of the 32 phenotyped novel ROS TFs we found at least one mutant allele with significant oxidative stress-induced changes in rosette area relative to the wild type (i.e. genotype effect dependent on the oxidative stress treatment and/or genotype-by-treatment effect under at least one of the tested stress treatments; Figure 4; Table 2). For the other novel TFs, no significant changes were detected by the genotype under all tested conditions, either there was an overall genotype effect under all tested conditions (similar changes were observed) or the genotype effect was independent of the stress treatments (e.g. WRKY47; Figure 4). For SCL13, WRKY45, BEH2 and PHL1, experimental evidence is provided by at least two mutant alleles. *WRKY45* loss-of-function (T-DNA mutant RATM11− 0634− 1_H and RNAi line AS3.1) affected the rosette growth during low and high 3-AT stress compared to the wild type in a genotype-by-treatment dependent matter, whereas, *WRKY45* OE (line 15.1) caused an increased rosette yield that was dependent on stress treatments low 3-AT and low MV (Figure 5). Two *SCL13* overexpression alleles (1.11 and 2.12) displayed decreased rosette growth dependent on both MV conditions in a genotype-by-treatment-dependent manner (Figure 5). Similarly, knockdown of *SCL13* (RNAi lines AS112D and AS28C) negatively affected rosette growth under MV stress, but no genotype-by-treatment interaction effect was observed. *PHL1* T-DNA and *PHL1/PHR1* double T-DNA lines displayed altered growth on low and high 3-AT, respectively, the latter dependent on the genotype-by-treatment. *BEH2* mutation altered the rosette area that was dependent on the stress treatments high 3-AT and low MV for line SALK_079430 and dependent on the genotype-by-treatment for line SAIL_76B06.

**Figure 4.**
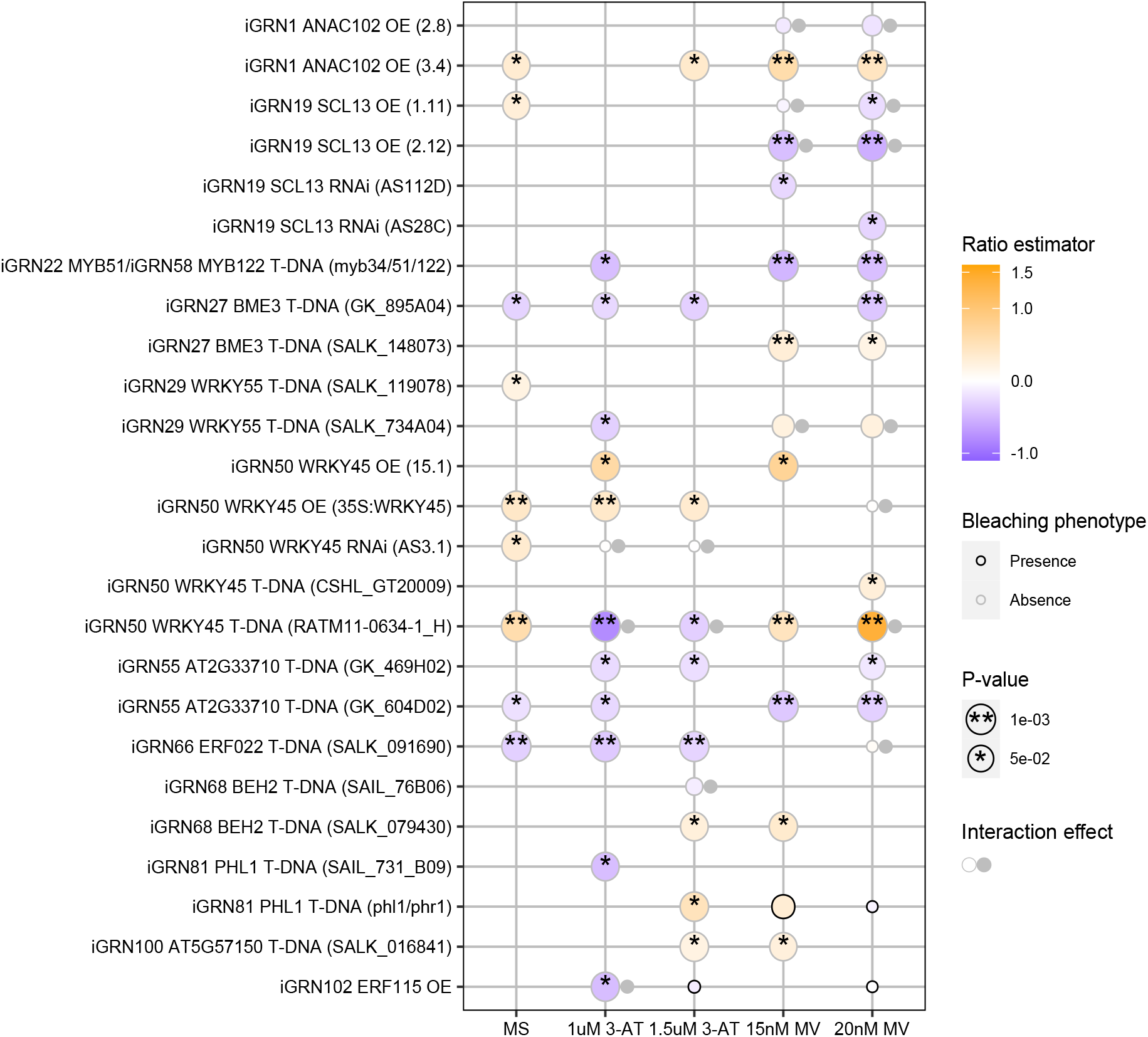
Oxidative stress phenotyping results of novel ROS TFs. Heatmap presenting phenotypes associated with mutant alleles of iGRN extracted novel ROS TFs under different oxidative stress conditions. Mutant lines with a statistically significant phenotype under oxidative stress (1 µM and 1.5 µM 3-AT and 15 and 20 nM MV) and/or normal growth conditions (MS) are displayed. Results of other mutant lines that were tested in this study can be found in Supplemental Table S5 and S6. Statistically significant differences relative to wild type (Col-0) are indicated by asterisk; *, *P* < 0.05 and **, *P* < 0.001 as determined by the hypothesis tests on the fixed effects of the mixed model (two-sided, see Methods). Data were obtained from at least two biological replicates per experiment and from independent experiments. Color scale indicates the ratio of the rosette area of 14-day-old mutant alleles relative to wild type (Col-0). Black and grey borders indicate presence or absence of a bleaching phenotype of the mutant plants, respectively. Small grey circle indicates a statistically significant genotype-by-stress treatment interaction effect.

**Figure 5.**
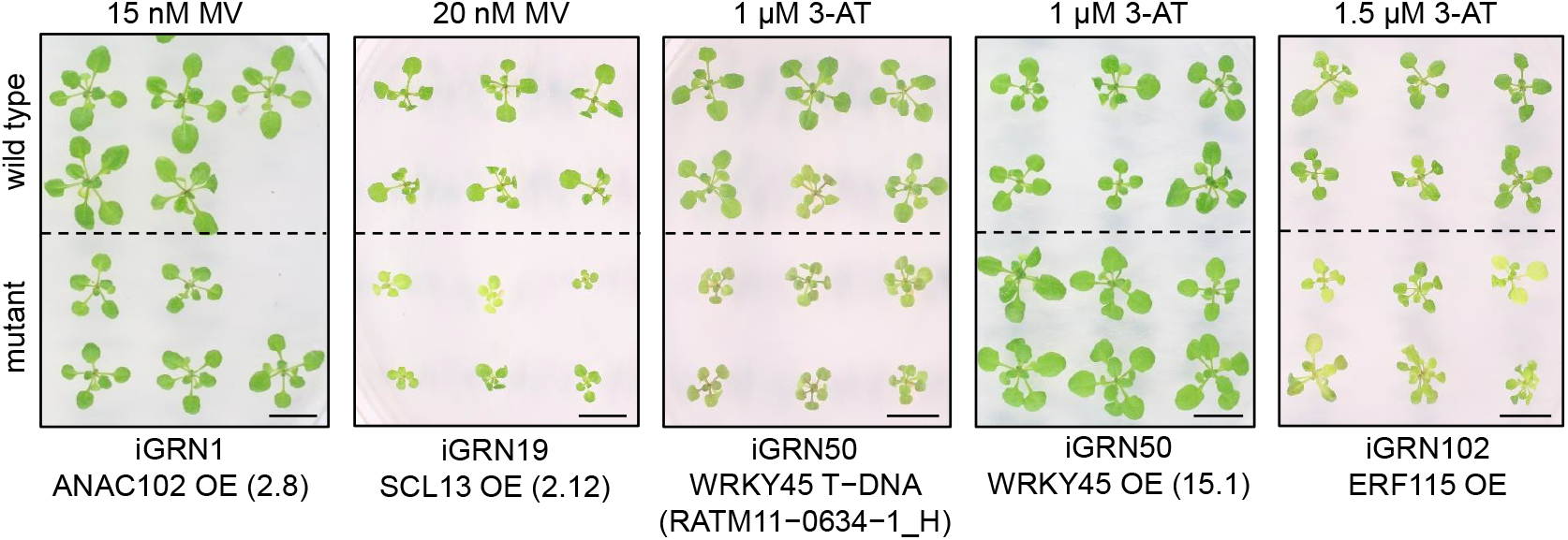
Representative images of oxidative stress phenotypes for a selection of mutants. Representative images of novel ROS TFs with a phenotype during oxidative stress. 17-days-old seedlings of selected mutant lines compared to wild type (Col-0). Scale bars, 1 cm.

We further demonstrated that ANAC102, BME3, MYB51, WRKY55, AT2G33710, ERF022, AT5G57150 and ERF115 have a function in oxidative stress responses based on one mutant allele (for *ERF022* and *ERF115* only one line was available) (Figure 4 and Table 2). For example, *ANAC102* OE line 3.4 (strong OE; ˜ 100-fold) displayed a larger rosette area that was independent of the stress treatment, while rosette area changes by *ANAC102* OE2.8 (weak OE; ˜4.5-fold) were dependent on the genotype-treatment interaction (Figure 5). *ERF115* OE affected plant performance in a manner that was dependent on the stress treatment, with a decreased rosette area under low 3-AT (dependent on the genotype-by-stress treatment) and severe bleaching under high MV and 3-AT concentrations (Figure 5).

Apart from validating the identified novel ROS TFs based on an oxidative stress-related growth phenotype, we evaluated the correct prediction of ROS responsive target genes. Therefore, we compared the predicted target genes of the highest ranked novel ROS TF ANAC102 with genome-wide target genes identified by ChIP-Seq^27^. Among the 543 ANAC102 target genes predicted by the iGRN, 185 (34%) were confirmed by ChIP-Seq, indicating a significant overlap (2.28 fold enrichment, p-value<2e-26). Meta-analysis of transcript profiles during various oxidative stress conditions shows that 103 of the 185 confirmed target genes are responsive to a wide range of ROS-inducing conditions^32^(Supplemental Table S8, Supplemental Figure S5) and include 20 out of the 35 ROS marker genes predicted to be ANAC102 targets. Among the 543 predicted ANAC102 target genes are also 62 TFs of which 28 are confirmed by ChIP-Seq. Among these ChIP-confirmed TFs are the ROS marker genes *WRKY40, ZAT6, ZAT10, ZAT12* and *ERF5*, indicating extensive TF-TF regulation of ROS responses downstream of *ANAC102*. Furthermore, to be a true regulatory interaction, the spatial-temporal expression characteristics of the TF and its target genes must be similar. To assess this, we explored transcriptome profiles of different rosette and root tissues from Genevestivator^40^to compare tissue-specific expression profiles of *ANAC102* with its ChIP-Seq-confirmed target genes. We observed common expression signatures, namely *ANAC102* and its target genes being expressed in juvenile, adult and senescent leaves as well as in different primary root zones and in the lateral root (Supplemental Figure S5). Taken together, using ANAC102 as a test case, we confirmed the prediction of true target genes, and consistency in oxidative stress responsiveness and spatial-temporal (basal) expression profiles between *ANAC102* and its targets.

Finally, the expression profiles of KNOWN_ROS and validated novel ROS TFs during different oxidative stress conditions were analyzed and compared ^32^. Fourteen out of the 17 known ROS TFs were identified as responsive to a wide range of oxidative stress conditions (|log_2_ fold change| > 1 and FDR < 0.01; Figure 6; Supplemental Table S7). Among the 13 newly validated novel ROS TFs, only half were ROS responsive at the transcript level. *WRKY55, MYB122* (for which the triple mutant *myb34/51/122* shows a phenotype, but not the *myb122* single mutant), *ERF022, BEH2, PHL1, AT5G57150* and *ERF115* expression levels are not regulated by oxidative stress, or only regulated in a small number or specific set of oxidative stress-related conditions, and thus, would not have been predicted from studies solely relying on differential expression at the whole plant or organ level. For example, *ERF115* has tissue-specific expression patterns, confined to the stem cell niche within the root meristem ^41^, indicating that transcript analysis on the whole plant level might not reveal stress-induced alterations in this specific tissue ^42^. Interestingly, we observed that novel ROS TFs showing a phenotype under oxidative stress have a 2-fold higher degree and 5-fold higher betweenness centrality compared to novel ROS TFs lacking such phenotype (median degree of 1670 and 749, median betweenness centrality of 699,318 and 142,777, for novel ROS TFs with and without oxidative stress phenotype, respectively).

**Figure 6.**
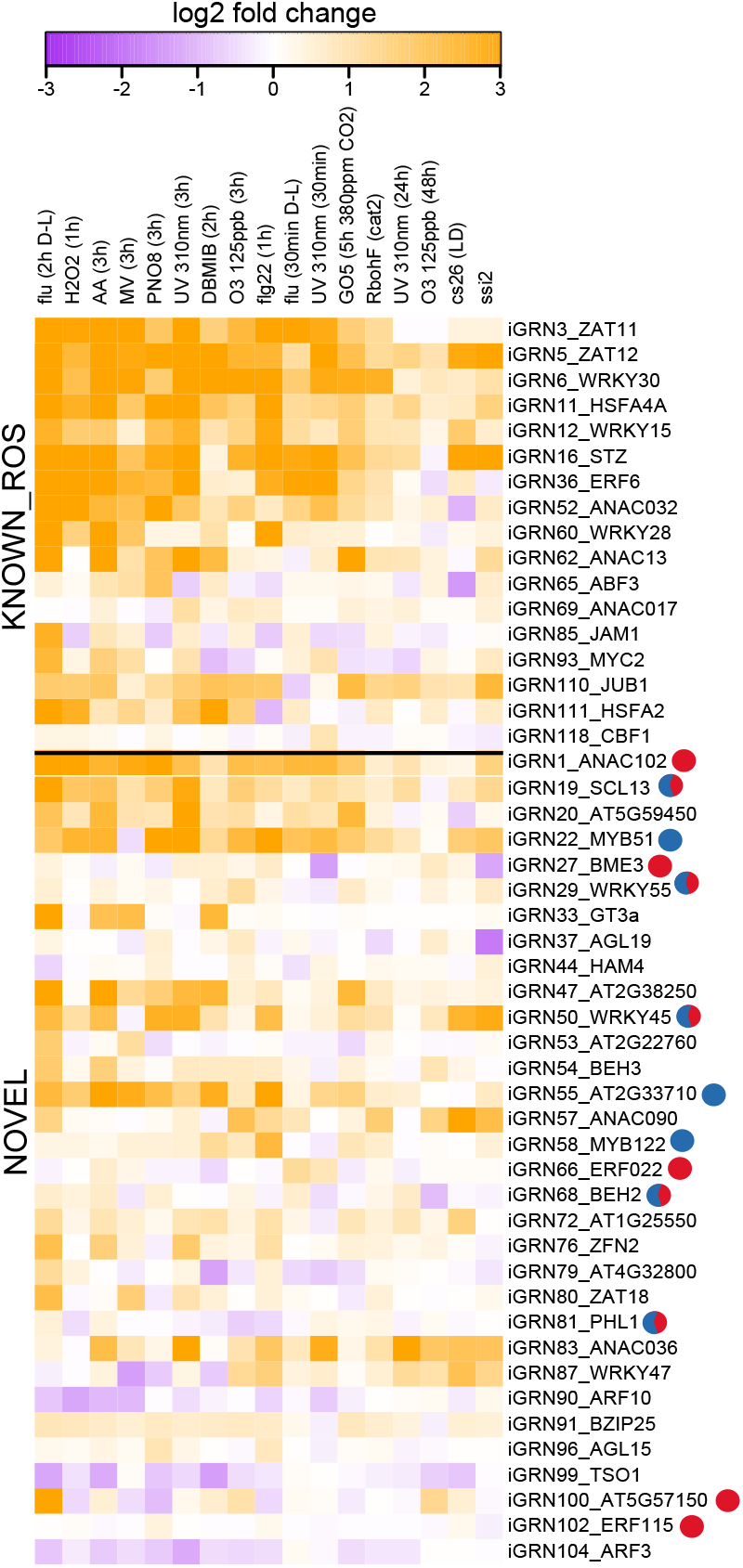
Expression of KNOWN and NOVEL ROS TFs during various conditions provoking cellular ROS/redox imbalances. Heatmap showing the expression response of ROS TFs extracted from the iGRN during a selection of conditions provoking cellular ROS production or redox imbalances (a more elaborate set of oxidative stress conditions is displayed in Supplemental Table S7). TFs with previously known function in ROS responses and TFs with no functional link with any oxidative stress and/or (a)biotic stress responses (NOVEL) that were tested in our phenotyping assays are displayed. NOVEL TFs validated in our study are shown using blue and red circles, indicating a positive and negative effect by the TF on oxidative stress tolerance, respectively.

Overall, among the 124 novel ROS TFs extracted from the iGRN, 75 (60%) were validated by previously published studies (before the iGRN was constructed) to be involved in the plant’s responses to ROS perturbation and/or (a)biotic stresses. By analyzing mutant lines perturbed in one of the analyzed 32 novel ROS TFs, 13 additional regulators were validated for a function in ROS responses in our phenotyping assays. Among the novel ROS TFs newly identified by our phenotyping assays, WRKY55, WRKY47, AT2G33710 [ERF] and AT5G57150 [bHLH] had previously not been associated with any function. Other validated novel ROS TFs are known to function in high-light stress tolerance or acclimation responses (NAC102, BME3 [GATA family] ^33-35^), age-triggered senescence (WRKY45, ERF022 [AP2/EREBP]; ^43,44^), response to phosphate starvation (WRKY45, PHL1 [G2 family]; ^45,46^), submergence stress (WRKY45; ^47^), glucosinolate (antimicrobial compounds involved in plant immunity) biosynthesis (MYB51, MYB122; ^48^), brassinosteroid signaling (BEH2 [BES1 TF family]; ^49^), phytochrome-dependent light signaling (SCL13 [GRAS family]; ^50^), hypocotyl growth (BEH2 [BES1 family]; ^51^), seed germination (BME3; ^52^) and somatic embryogenesis (ERF022; ^53^). Overall, among the novel ROS TFs validated for an oxidative stress function by literature evidence or in our phenotyping assays, the AP2-EREBP (5), NAC (5) and WRKY (5) families are highly represented (Supplemental Table S5). These families are typically associated with abiotic, biotic and/or oxidative stress responses ^54^and the iGRN identified additional members with an oxidative stress phenotype. In addition, the iGRN correctly predicted an oxidative stress response function for one or more members of the bHLH, ZAT (or C2H2), HSF, MYB and bZIP families that are also often associated with functions in plant stress responses. The iGRN also identified novel ROS TFs from families that were known to be mainly involved in plant growth/developmental processes, such as GRAS (SCL13), BES1 (BEH2) and GATA (BME3) (^49,55,56^). So far there is very little evidence for the involvement of GATA TFs in plant stress responses except for reported functions for rice and soybean GATA TFs in low nitrogen stress tolerance ^57,58^, the identification of salt and drought stress-induced alternative splice products for rice GATA TFs ^59^, and the recent characterization of BME3 in systemic acquired acclimation to high light stress, downstream of a systemic signaling cascade mediated by ROS signaling ^34,35^. Consistent with recently published studies ^34,35^, we show that BME3 is one of the first GATA transcription factors with a proven function in the regulation of stress responses.

## DISCUSSION

The comprehensive experimental mapping of plant GRNs is a major challenge and different technologies, either *in vitro* or *in vivo*, each provide partial snapshots of the regulatory logic in *Arabidopsis*. While different integrative approaches have been applied to identify regulatory networks controlling specific biological processes in plants ^12,60-63^, in this study we leveraged a wide variety of regulatory datasets capturing different properties of TF-mediated transcriptional control. Through the application of an advanced machine learning approach for large-scale functional data integration, we performed for the first time a systematic regulatory annotation for nearly all *Arabidopsis* genes. Although the presented iGRN is generic and not tailored towards specific plant organs or growth stages, extensive validation using different types of experimental datasets, measuring TF binding and/or regulation, revealed a strong enrichment for functional interactions. In addition, the iGRN outperformed all input networks when assessing the predictive power using a gold standard evaluation set, demonstrating the success of the supervised learning approach. An assessment of the functional coherence of a TF’s target genes suggests that the iGRN regulatory interactions are of equally high quality compared to those obtained through experimental methods. Taken together, our validation results indicate that the iGRN is strongly enriched for functional interactions and that the applied integrative approach clearly complements GRNs inferred through a single experimental assay.

Investigating the relationship between network architecture and the complexity of TF control confirmed that TFs themselves are complexly regulated by other TFs. In addition, through the integration of large-scale phenotyping data, a significant association was found between a TF’s network centrality (but not degree) and observing a phenotype when knocking out that TF. A similar pattern was observed for the novel ROS TFs showing an oxidative stress phenotype. This finding indicates that these TFs occupy key positions in the GRN and loss of their gene products has major consequences for plant growth and development. Furthermore, TFs showing a phenotype when screening mutant lines were not having more TFs among their target genes compared to TFs lacking phenotypes, suggesting that master regulators high in the TF network hierarchy are not more frequently showing phenotypes compared to lower-tier TFs. Obviously, these findings depend on the global quality of the iGRN but complement previous results reporting that higher-tier TFs in ABA networks do not show more differential expression during ABA response ^27^. Finally, TF pairs sharing many target genes are indicative of combinatorial control, where TFs physically interacting with each other form higher-order protein complexes mediating increased DNA-binding specificity ^64^. Consequently, TF pairs with many shared target genes in our integrative network which also show correlated expression profiles are good candidates for future research to shed light on the role of cooperative TF control in specific biological processes in plants.

Predicted TF functions based on the iGRN regulatory interactions indicated that for various biological processes many known regulators could be recovered. Starting from our systematic regulatory annotation, known functional annotations were confirmed for 681 TFs. For several biological processes our predicted TF functions are highly complementary to the AraNet probabilistic functional gene network and for 268 TFs lacking experimental annotations, new functions were inferred. To validate the power of our integrative network, novel regulators predicted to play a role in oxidative stress signaling were experimentally validated using ROS-specific phenotyping of loss- or gain-of-function lines. By analyzing transgenic lines perturbed in one of the 32 novel ROS TFs, an additional 13 (41%) regulators were validated for a function in ROS responses by our phenotyping studies. In addition to TF families that have often been associated with oxidative or other environmental stress responses, the iGRN identified novel ROS TFs from the GRAS, BES1 and GATA families that are not, or to a lesser extent, known to be involved in stress responses. Notable, compared to previous studies that predicted gene functions or prioritized candidate regulators based on (co-)expression analysis ^65^, our method enables the identification of novel regulators that would have been missed when solely relying on (differential) expression information. To conclude, our integrative network offers a high-quality resource to study TF control for individual target genes and to identify, through the interrogation of specific or condition-specific gene sets, novel regulators controlling different plant traits.

## METHODS

### Input datasets

We assembled a comprehensive compendium of genome-wide datasets that were used as primary data sources to construct the seven input networks and to build the integrative gene regulatory network. Starting from the TAIR10 Arabidopsis gene annotation, we added a set of 5711 non-coding RNAs described in ^66^resulting in a dataset covering 38,966 genes. A first dataset consisted of 1,260 PWM binding site profiles for 874 transcription factors. This dataset contained 108 and 623 positional frequency matrices that were obtained from protein binding microarray studies performed by ^67^and ^68^, respectively, and 529 positional frequency matrices obtained from DAP-seq experiments ^69^. A second dataset contained all available DNaseI hypersensitive data obtained from the plant regulome data resource (http://www.plantregulome.org/, ^14^). All separate conditions were merged into a single open chromatin dataset. A third dataset, composed of evolutionary conserved binding sites for 874 TFs, was obtained using the comparative motif mapping algorithm as described ^15^, starting from the input motif dataset as described above.

A fourth dataset comprised gene expression profiles generated by RNA-seq. The RNA-Seq expression compendium was built with public datasets from NCBI’s Sequence Read Archive (SRA) using Curse ^70^. The compendium contains gene-level expression values for 40 manually selected samples (Supplemental Table S9) of different treatment and tissue combinations (in total >2,7 billion reads). SRA files for each sequencing run were downloaded from the SRA and converted to the FASTQ format using fastq-dump (v2.4.4) from the SRA toolkit. FASTQ files from runs of the same sample were concatenated. Paired-end reads were unpaired by randomly selecting either the forward or reverse read and processing it as single-end. FastQC (v0.9.1) was used to detect overrepresented adapter sequences, which were subsequently clipped with fastx_clipper from the FASTX toolkit (v0.0.13). Nucleotides with Phred quality scores lower than 20 were trimmed with fastq_quality_trimmer from the FASTX toolkit. Reads shorter than 20 nucleotides after quality trimming were discarded. To obtain raw read counts for each transcript in the TAIR10 annotation, Sailfish (v0.6.3) ^71^was run with a k-mer length of 20. For genes with multiple transcripts, the raw read counts of its transcripts were summed to get a gene-level read count. Counts were then normalized for the entire compendium with the Variance Stabilizing Transformation (VST) from the DESeq R package (v1.14.0) ^72^. VST was chosen since it results in correlation coefficients between genes that are most comparable to those obtained with microarray data ^73^. A fifth dataset consisted of a compendium of 27 genome-wide TF binding datasets obtained through ChIP-seq ^7^.

### Network inference for the different input datasets

We derived input networks from each dataset and a weight was assigned to each TF-target gene interaction reflecting the certainty of the regulatory interaction. The motif target gene network (PWM) was created by mapping all binding site data on an extended locus for each gene in the *Arabidopsis thaliana* genome spanning 2kb upstream, intron and 1kb downstream regions, with the exons masked. All binding site specificity matrices were mapped genome-wide using Cluster-Buster with C parameter set to zero and M parameter set to 5 ^74^. Both the C and M scores were used as weights, which refer to the cluster and motif score of Cluster-Buster (Supplemental Table S3). Due to the short length of motifs, many instances of motifs in a single genome are expected to be coincidental matches that may not be bound *in vivo* or have no functional regulatory effect. Therefore, the resulting set of binding sites was filtered using DNaseI hypersensitive sites ^14^in order to obtain an open chromatin filtered motif network (PWM in DHS). The signalValue reported by the authors was retained as weight for motif matches in open chromatin, this is a measurement of overall enrichment for the region to DNaseI hypersensitivity. A second filtering approach was based on the evolutionarily conservation of these binding sites resulting in a conserved motif network (Conserved PWM). Weights in the conserved motif network are defined by the species conservation score according to the Comparative Motif Mapping (CMM) method as described in ^23^. Co-expression clusters were built around all TFs obtained from PlantTFDB ^75^using the Pearson correlation coefficient and only the top 1000 most co-expressed genes were retained as putative target genes (CoE TF-target). GENIE3 was run using standard parameters with the same list of regulators, only predicted interactions with a variance in expression explained larger than 0.005 were retained (GENIE3) ^16^. A k-nearest neighbor (kNN) co-expression cluster of size 100 was built around each gene in the *Arabidopsis thaliana* genome using the RNA-seq compendium as defined above.

PWM motif enrichment was performed on this co-expression cluster using the hypergeometric distribution and significantly enriched motifs (p-value < 0.001 after Benjamini–Hochberg correction for multiple hypotheses testing) were assigned to the members of the co-expression cluster in order to obtain the kNN motif enrichment network (CoE + PWM). The enrichment fold of the motif in the co-expression cluster is used as a weight. The peak scores for each TF bound region assigned to the closest gene were used as weight in the ChIP-seq network (TF ChIP) ^7^.

### Integrative network inference

The integrative network inference method is based on a binary classifier, where the class label represents the presence or absence of an interaction. The set of true interactions (positives) used in the training set was obtained from all validated interactions in AtRegNet ^18^, a review article on known interactions in secondary cell wall development ^19^and large scale data mining effort on 974 peer reviewed articles called ATRM ^20^, and has 5,732 edges, connecting 522 TFs with 3,738 targets. The remaining interactions between these TFs and potential target genes are considered to be negatives. In other words, while no true “negatives” are present in the dataset, we assign negative labels to TF–target pairs where the TF appears in the TFs of one of the positive pairs and this TF-target pair is not in the positive pairs. The negative labels reflect the fact that the TF has been studied and since no interaction is reported for it, it is more likely to represent a negative interaction. This dataset was split into a dataset for learning (80%) and a dataset for independent evaluation (20%). As such, the training set consists of 1,951,236 TF– target gene pairs, out of which 4,586 are positives (true interactions) and 1,946,650 negatives. The remaining gene pairs for TFs absent from the learning/evaluation dataset are considered to be negatives. Using the weights derived from the input networks we tested a number of different classification methods (gene pairs missing in specific input networks received a weight of zero in the matrix used for learning). We extracted the 4,586 validated interacting pairs from the true dataset and obtained the negative pairs for training by selecting a random subset of 13,758 (three times the number of positives; 3x 4,586) negatively labeled interacting pairs. Using this labeled training set different classifiers were built using the weights of the input networks as features. All features were normalized to have zero mean and unit variance. Classifiers were evaluated using the 20% evaluation dataset. In this analysis the gradient boosting using decision trees, random forests, and logistic regression were compared. The gradient boosting model made use of 1,000 trees, shrinkage of 0.01 and an interaction depth of 3. The random forest model used 500 trees selecting a random set of features of size square root (features) was sampled in each iteration. Logistic regression was used with a binomial distribution suitable for binary classification. All classifiers were compared using the area under the curve (Supplemental Table S2).

The final network was generated using the gradient boosting classifier, as this was the best performing throughout the training and validation steps. A total set of 60,714,507 unlabeled interactions was classified. We included all interactions that are predicted to interact with a probability of 0.35 or higher in the integrative network as this cutoff maximized the F1 measure on the test dataset (Supplemental Table S2). The full iGRN can be mined using overlap and TF enrichment analysis at http://bioinformatics.psb.ugent.be/webtools/iGRN/. Code, datasets and documentation describing the supervised learning approach are available at https://github.com/VIB-PSB/iGRN.

The overlap between the iGRN and the evaluation dataset was evaluated by counting how many interactions from these experimental interactions were present in the integrative network. The enrichment between two networks was defined as the number of interactions that are present in both networks divided by the number of interactions expected by chance. The number of common interactions expected by chance is given by the mean overlap of 10,000 randomized networks ^21^. Network randomization was done by permuting the labels of all TFs and the labels of all genes, which preserves the network structure.

### Functional evaluation using experimental datasets

Three experimentally obtained gene regulatory networks were used for further validation of the predicted network. An accumulation of publicly available Y1H data from literature was used to generate a Y1H network containing 2,759 interactions ^76-79^. A set of differentially expressed genes upon TF perturbation was obtained from the CORNET database, containing 51,178 interactions ^80^. An independent set of interactions determined by ChIP-seq was obtained from literature, reporting 148,949 interactions ^27,81-84^. From Song et al. 2016 only non-ABA conditions were considered.

The overlap between the iGRN and three experimentally determined gene regulatory networks was evaluated by counting how many interactions from the experimental networks were present in the integrative network. The enrichment between two networks was defined as the number of interactions that are present in both networks divided by the number of interactions expected by chance, as explained above. Network randomization was done by permuting the labels of all TFs and the labels of all genes, which preserves the network structure and assures that the observed enrichment is not due to potential biases arising from structural properties of the network (Supplemental Table S3).

Functional enrichment was determined for each sub network (intersection, FN, FP) using biological metrics as described in ^23^using Gene Ontology information. Note that no functional GO information was used as input for the iGRN construction.

### Analysis of network architecture and overlap with plant phenotypes

Based on the target genes for all TFs present in the iGRN, degree and betweenness centrality were computed using the igraph R package, considering directed edge information. Three levels of network hierarchy were calculated using the R code available from http://archive.gersteinlab.org/proj/hinet/ ^29^. The Jaccard index, defined as the intersection over union, was used to compute the overlap of target genes between two TFs. Based on bulk downloads of the RARGE II database (^28^; download November 2018), which reports ontology-based phenotype information for 48,077 lines (17,671 RIKEN Arabidopsis Ds transposon mutant lines, 16,337 CSHL Arabidopsis genetrap mutant lines, 14,069 RIKEN Arabidopsis full-length cDNA overexpressed Arabidopsis lines). The presence of phenotypes in the RARGE II dataset was scored for different sets of regulators, defined based on degree and betweenness centrality.

### TF function prediction

For each TF, iGRN target genes were subjected to GO enrichment analysis using the hypergeometric distribution combined with Benjamini–Hochberg (B&H) correction for multiple hypotheses testing. A q-value cutoff 1e-03 was applied since this resulted in the best F1 score, a metric that combines precision and recall to recover known GO annotations. Only experimental and curated GO BP annotations were considered (version August 2018), GO annotations were propagated to include parental GO terms, and a set of 18 general GO terms, corresponding with terms located at the root of the GO hierarchy, were excluded (GO:0008150, GO:0009987, GO:0008152, GO:0044237, GO:0071704, GO:0050896, GO:0065007, GO:0032502, GO:0050789, GO:0032501, GO:0007275, GO:0050794, GO:0006355, GO:0045893, GO:0045892).

To predict ROS TFs, core genes that are part of the ROS wheel clusters IV-VIII, covering genes related to chemical and genetic perturbation of cellular ROS/redox, were used to define a custom category of ROS marker genes ^32^. Subsequently, the over-representation of these ROS markers in the set of iGRN target genes per TF, was used to predict novel ROS TFs and rank them based on the q-value for enrichment (hypergeometric distribution, q-value cutoff 1e-05). The assignment of ROS TFs to the categories KNOWN_ROS, KNOWN_STRESS and NOVEL_TF was based on the information available in the literature at the moment the initial iGRN was constructed (December 2016).

### Phenotyping candidate ROS TFs

T-DNA insertion mutants were obtained from the European Arabidopsis Stock Centre (NASC; http://arabidopsis.info/), GABI-Kat (http://www.gabi-kat.de/db/), Versailles Arabidopsis Stock Center (http://publiclines.versailles.inra.fr/tdna/index) (Supplemental Table S6). Homozygous T-DNA insertion lines were selected by PCR analysis (Supplemental Table S10). Mutant lines previously published were provided by the authors (Supplemental Table S6 (overview lines)). Overexpression lines were generated by Gateway cloning in the binary destination vector pK7WG2D (18056864) (Supplemental Table S10). Plants were grown on half-strength Murashige and Skoog (1/2MS) medium (Duchefa) supplemented with 1% (w/v) sucrose, 0.75% (w/v) agar, and B5 vitamins, pH 5.7, at 21°C and 60 μE m^− 2^s^− 1^light intensity in a 16-h/8-h light/dark photoperiod. To induce oxidative stress, the 1/2MS medium was supplemented with 15 or 20 nM MV; or 1 or 1.5 μM 3-AT, causing on average 15% (15 nM MV), 30% (20 nM MV), 20% (1 µM 3-AT) and 55% (1.5 µM 3-AT) growth reduction compared to the nontreated seedlings. Rosette area was determined by automated area measurements of individual plant rosettes using ImageJ. Raw phenotyping results are available in Data file 3. Presence or absence of chlorosis/bleaching was monitored by visual assessment. For each mutant line, the rosette area of mutant seedlings was compared with wild-type plants grown in the same Petri plate. Data were obtained from at least two biological replicates per experiment and from independent experiments.

### Statistical analysis leaf phenotyping data ROS TFs

A mixed model was fitted to the rosette area with genotype and treatment as fixed effects as well as the interaction term with the mixed procedure from SAS (SAS Institute Inc. 2018. SAS/STAT® 15.1, Cary, NC). The factor plate was added as random effect in the model to take into account the correlation between observations made on the same plate. The Kenward-Rogers method was used to calculate the denominator degrees of freedom of the fixed effects. The null hypotheses of interest were an equal genotype effect upon treatment in the mutant line and the wild-type line and an equal treatment versus MS effect between the mutant line and the wild-type line (i.e. no interaction effect). A significance level of 0.05 was taken. To correct for multiple testing, the maxT procedure was used as implemented in the plm procedure.

## ACKNOWLEDGEMENTS

The authors wish to thank Cordelia Bolle (SCL13), Xuemei Chen (ARF3), Yi-Fang Chen (WRKY45), Donna Fernandez (AGL15), Henning Frerigmann (MYB51, MYB122), Małgorzata Gaj (ERF022), Tamara gigolashvili (MYB51), Dwayne Hegedus (SCL15), Lars Hennig (AGL19), Ildoo Hwang (BME3), Zhongchi Liu (TSO1), Lars Østergaard (ARF3), Javier Paz-Ares (PHL1) and Olivier Van Aken (WRKY45) for kindly providing Arabidopsis mutant lines. We thank Zoé Joly-Lopez and Olivia Wilkins for providing in-depth comments on an earlier version of the manuscript, as well as Camilla Ferrari and Indeewari Dissanayake for proofreading. I.D.C. is supported by the Research Foundation Flanders (postdoctoral fellowship 12N2415N and grant for a long stay abroad V400620N). J.V.d.V. is indebted to the Agency for Innovation by Science and Technology in Flanders for a predoctoral fellowship. X.L. and L.L. are supported by the China Scholarship Council for a PhD fellowship (201706910099 and 201808530499). F.V.B. is supported by the Fonds Wetenschappelijk Onderzoek – Vlaanderen (FWO) and the Fonds de la Recherche Scientifique – FNRS under EOS Project No. 30829584 and the Research Foundation Flanders (project nos. G0D7914N and G055416N).

## AUTHOR CONTRIBUTIONS STATEMENT

J.V.d.V., I.D.C. and K.V. designed the research. I.D.C., X.L., L.L. and R.P. performed the phenotype screening. V.S. performed statistical data analysis. D.V. generated the expression compendium. J.V.d.V. and K.V. performed network analysis and function prediction. J.V.d.V., I.D.C., F.V.B. and K.V. wrote the manuscript.

## OVERVIEW OF SUPPLEMENTAL FIGURES AND TABLES

**Figure S1. Overview of the iGRN interactions supported by different input networks. Figure S2. Overview of the in- and out-degree for the complete iGRN.**

**Figure S3. Examples of predicted cooperative TFs supported by protein-protein interactions.**

**Figure S4. Recovery of known GO annotations for TFs belonging to different biological processes.**

**Figure S5. Oxidative stress-responsive and tissue-specific expression profiles of ANAC102 and its ChIP-confirmed target genes.**

**Supplemental Table S1. Overview of TF families in gold standard.**

**Supplemental Table S2. Overview of data types, training methods and performance results for different machine learning classification algorithms.**

**Supplemental Table S3. Overlap and enrichment between different networks.**

**Supplemental Table S4. Overlap of target genes between iGRN and ChIP-Seq data.**

**Supplemental Table S5. Overview of all TFs predicted by the iGRN with a function in ROS responses.**

**Supplemental Table S6. Overview of mutant lines used for phenotypic analysis.**

**Supplemental Table S7. Expression of KNOWN_ROS and NOVEL ROS TFs during various conditions provoking cellular ROS/redox imbalances.**

**Supplemental Table S8. Expression of ANAC102 and its ChIP-confirmed target genes during various conditions provoking cellular ROS/redox imbalances.**

**Supplemental Table S9. Overview of samples included in the RNA-Seq expression compendium.**

**Supplemental Table S10. Primers used in this study.**

## DATA AVAILABILITY

Source data for Figs. 1,4 and S3 are provided with the paper.

**Data file 1. Data_file1_iGRN_supervised_network_support.txt All TF-target interactions in the iGRN**.

Columns report: TF, target genes, score, rank, support input networks.

Overview of generated, analysed and used iGRN regulatory interaction data for Fig. 1.

**Data file 2. TF_function_all.txt reporting predicted functions for all TFs present in the iGRN**.

The sheet README explains the different columns and offers 3 examples how this file can be explored for specific TFs, functions or a specific target gene.

Overview of generated, analysed and used iGRN TF function data for Fig. S3.

**Data file 3. Raw phenotype results that were used as input for the statistical analysis**.

Overview of raw phenotyping measurements for Fig. 4.

