## Supplemental figures for "Integrative inference of transcriptional networks in Arabidopsis yields novel ROS signalling regulators"

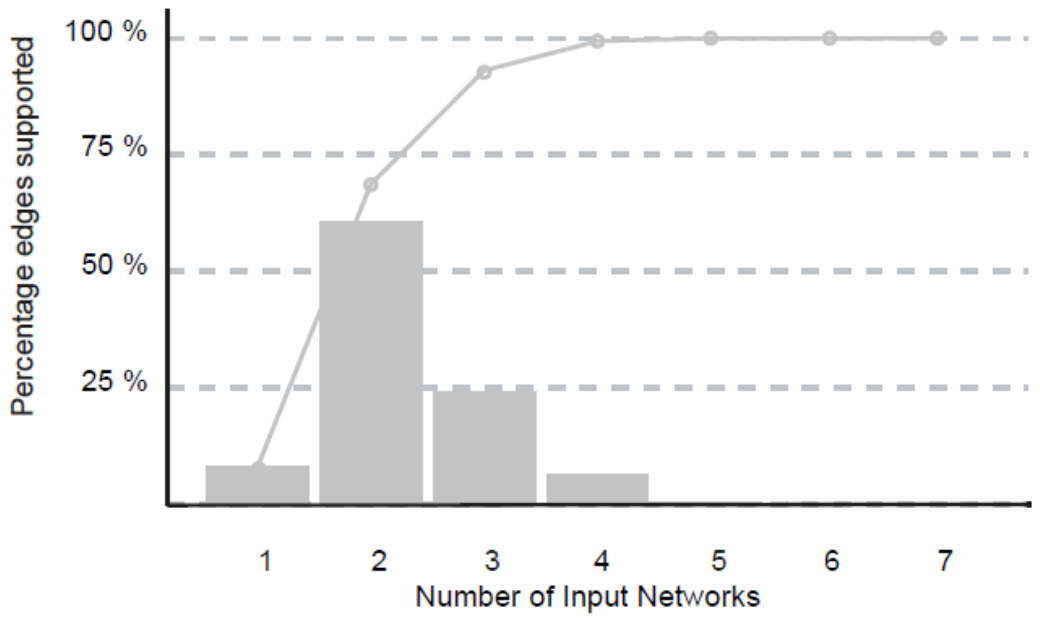

### Figure S1. Overview of the iGRN interactions supported by different input networks.

The grey bars report the fraction of interactions supported by the number of input networks, while the line shows the cumulative fraction.

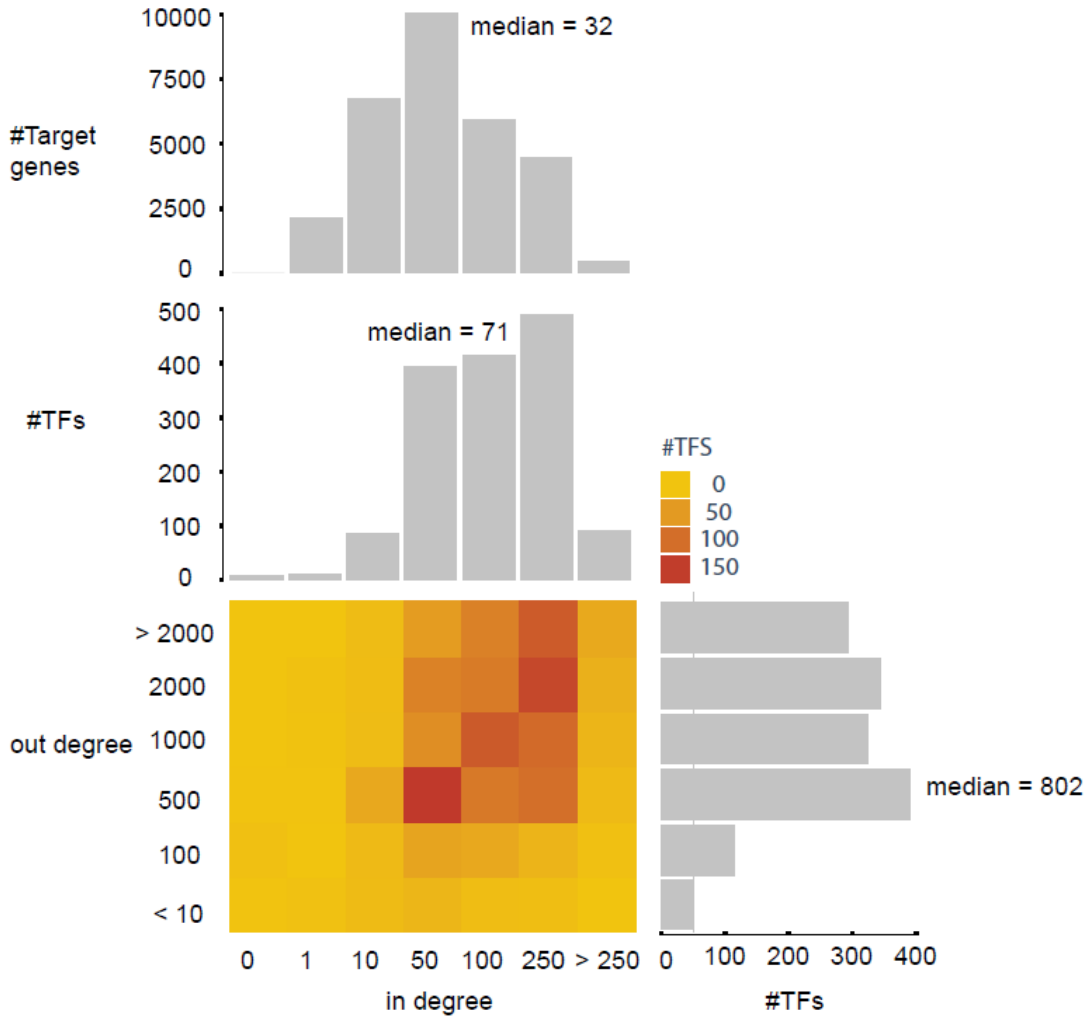

### Figure S2. Overview of the in- and out-degree for the complete iGRN.

The two upper bar plots, marked by the blue rectangle, show the in-degree (i.e. number of incoming TFs) for all target genes and TFs, respectively. The bar plot at the right, part of the green rectangle, shows the out-degree (i.e. number of target genes) for the different TFs (binned). The central bar chart depicts the in-degree for TFs in the network, showing the median number of incoming TFs is larger for TFs compared to considering all target genes (71 and 32, respectively). The heatmap shows the largest number of TFs have an in-degree of 50 incoming TFs and out-degree of 500 target genes.

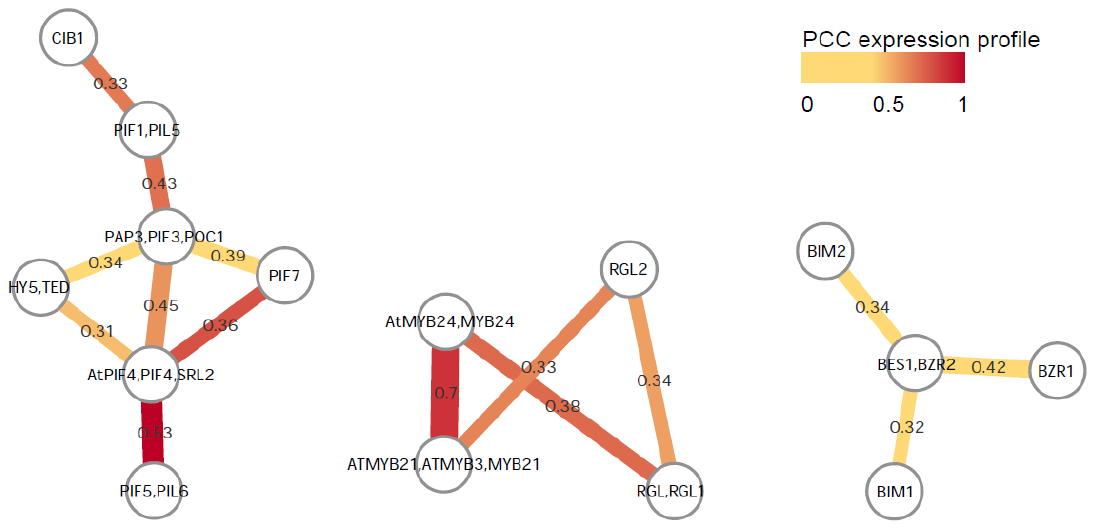

### Figure S3. Examples of predicted cooperative TFs supported by protein-protein interactions.

Sub-networks showing experimental protein-protein interactions for TFs sharing a large number of target genes (Jaccard index >0.3). Whereas the edge thickness and values reports the fraction of shared target genes for a pair of TFs quantified using the Jaccard index, the color represents the absolute value of the Pearson Correlation Coefficient between the expression profiles of a pair of TFs.

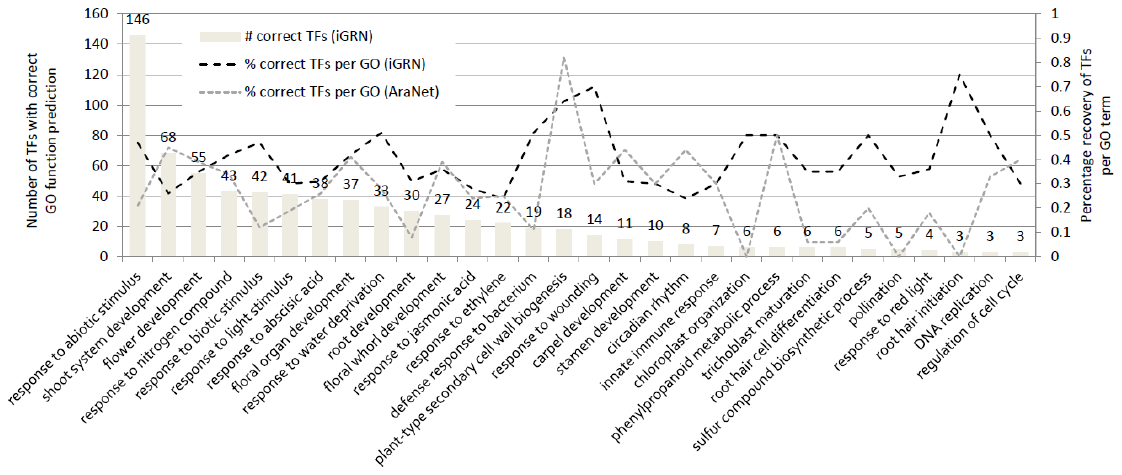

### Figure S4. Recovery of known GO annotations for TFs belonging to different biological processes.

The bar chart reports the absolute number of TFs with correctly inferred GO Biological Process annotations based on the iGRN. Dashed lines report the fraction of correctly predicted GO annotations for different Biological Processes based on iGRN and AraNet v2 (black and grey line, respectively).

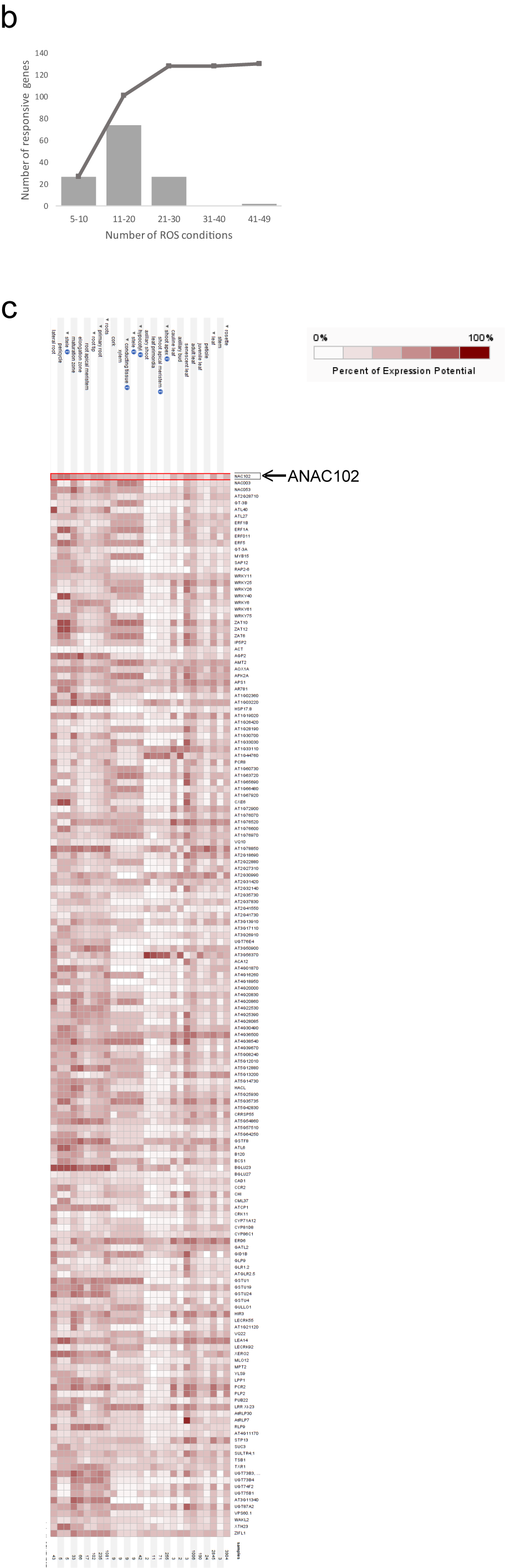

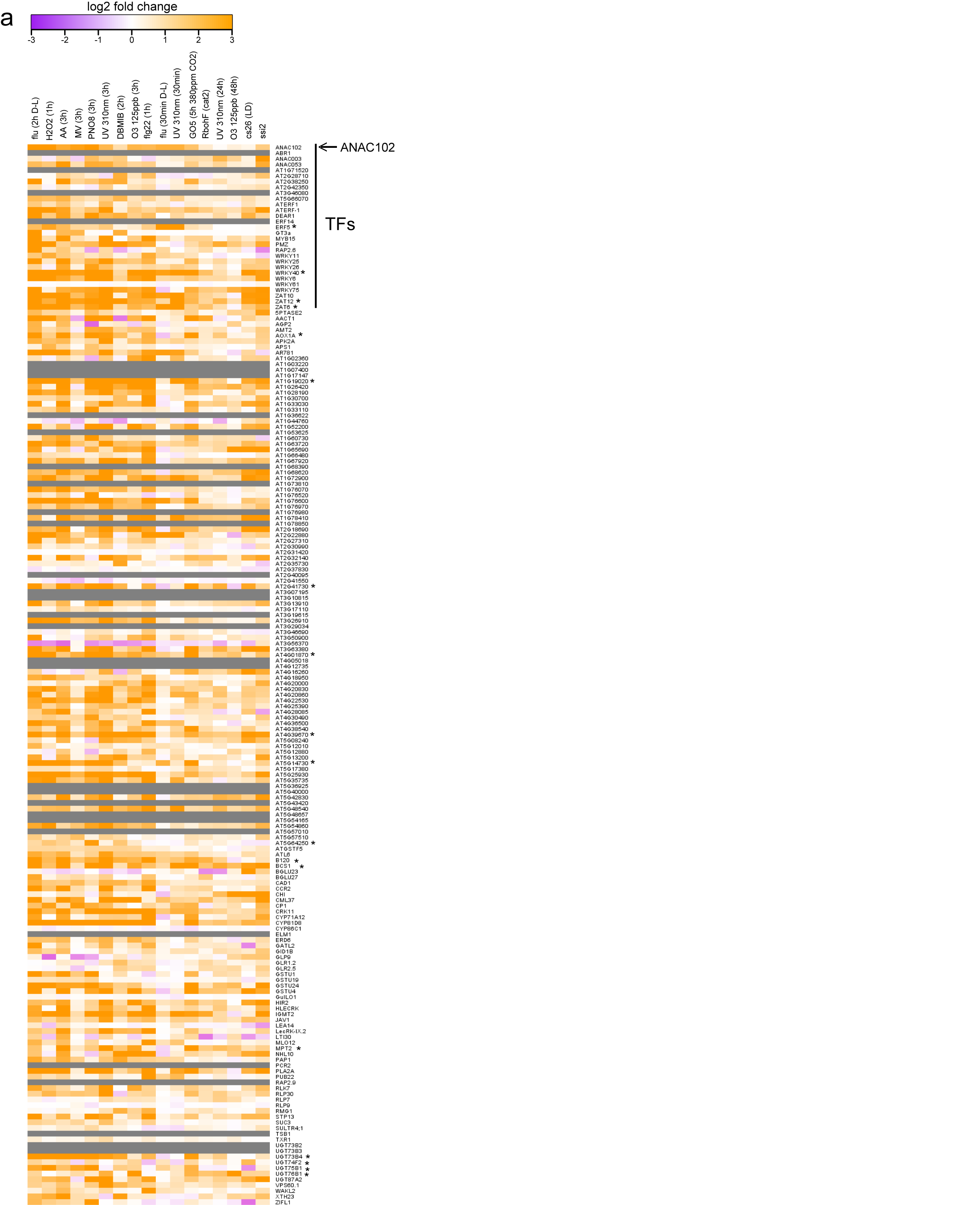

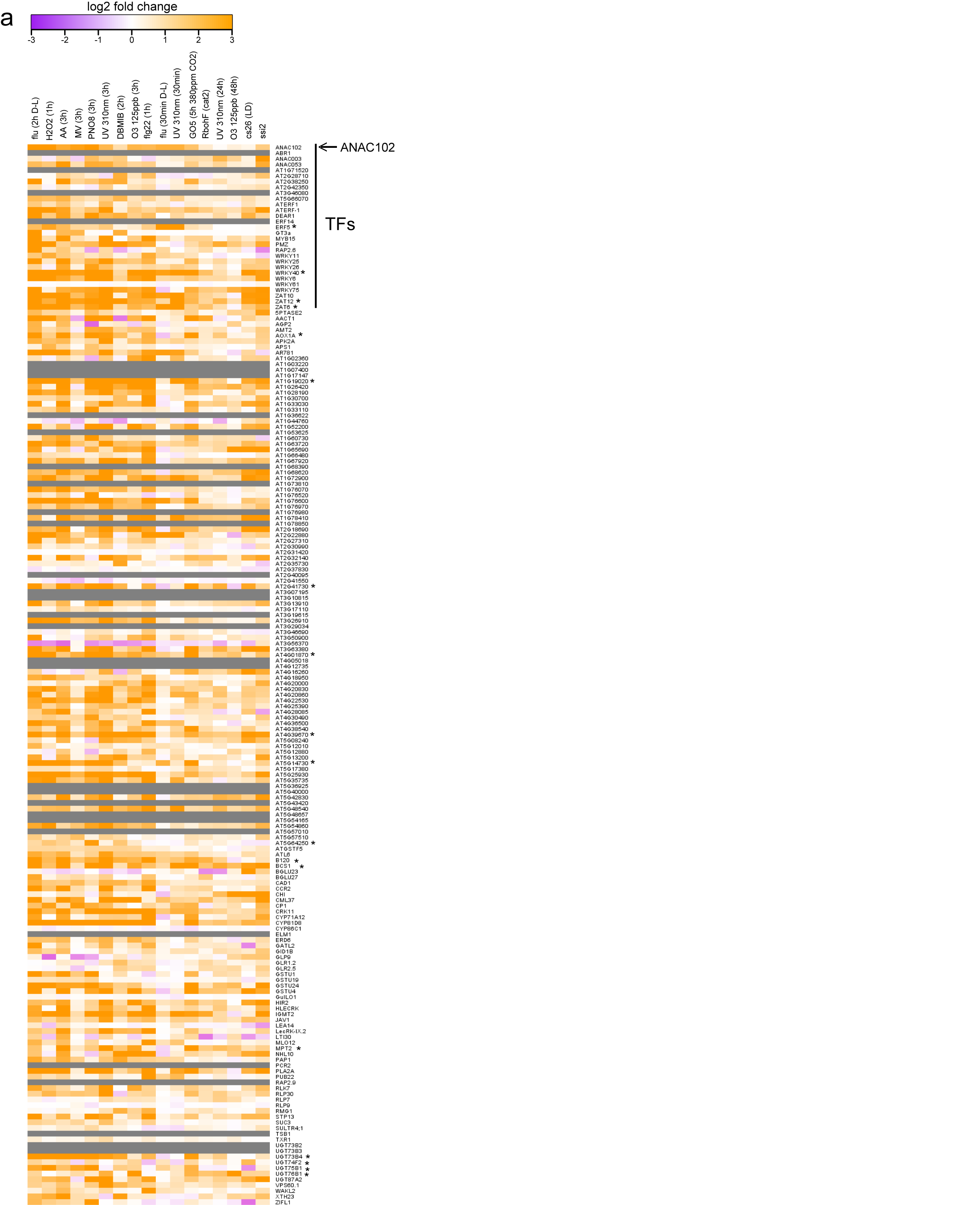

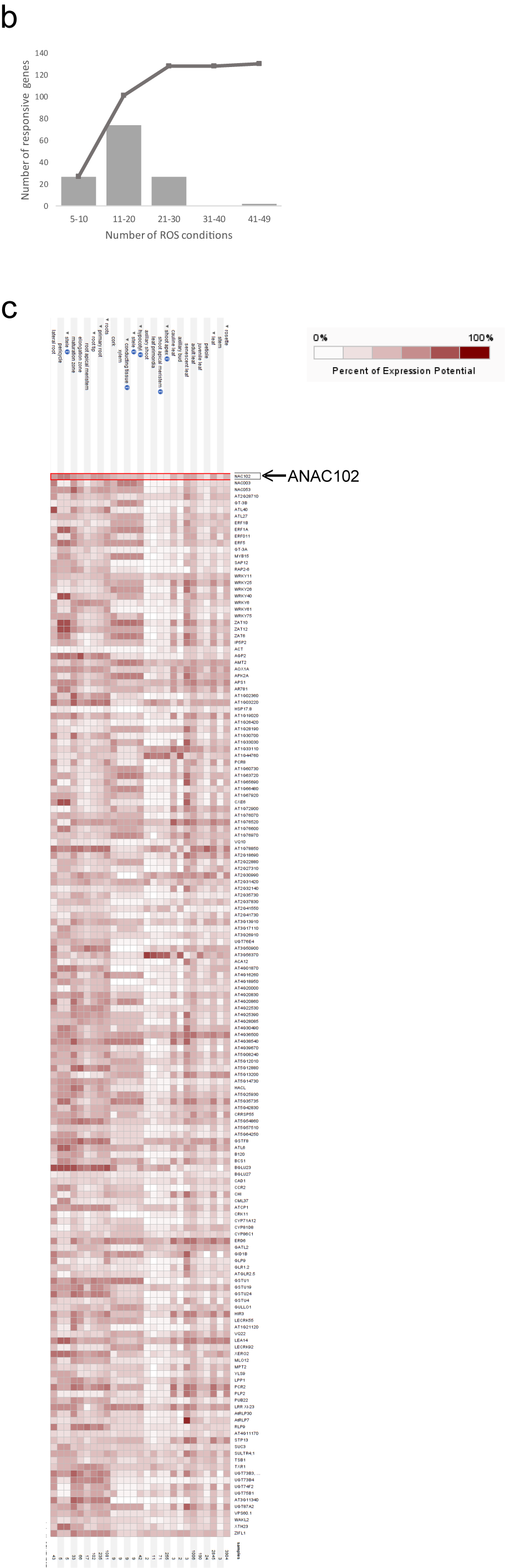

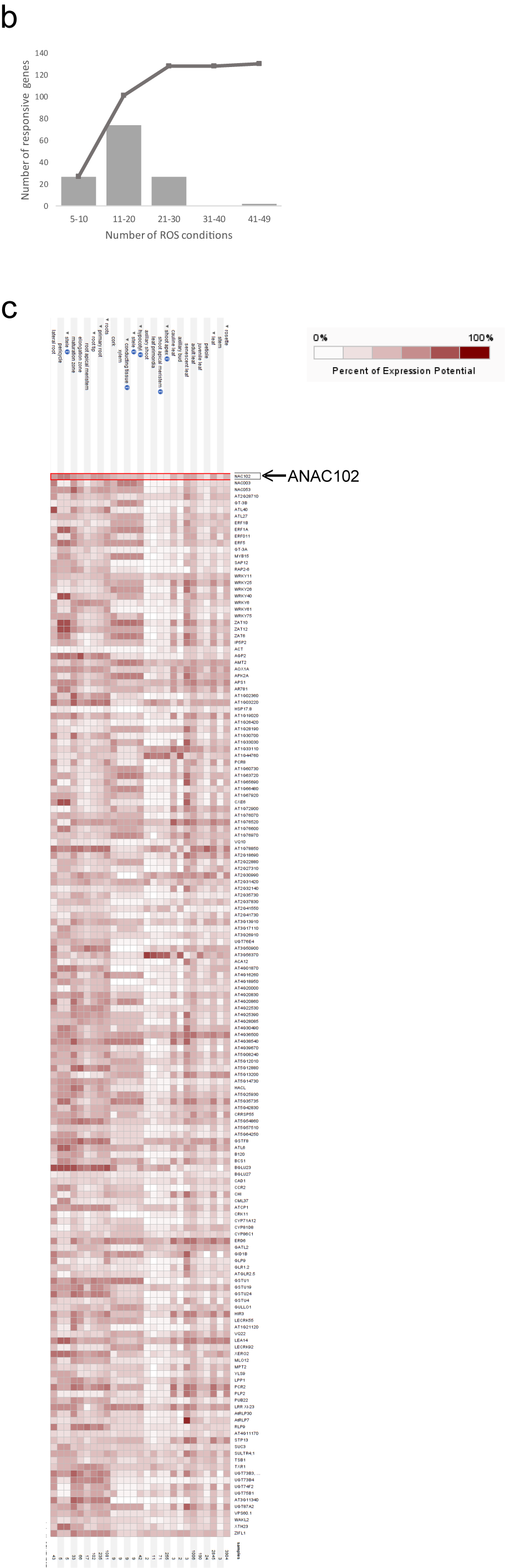

### Figure S5. Oxidative stress-responsive and tissue-specific expression profiles of ANAC102 and its ChIP-confirmed target genes.

A. Heatmap showing the expression response of *ANAC102* and its predicted target genes that were confirmed by ChIP during various conditions provoking oxidative stress (Willems et al., 2016) (a more elaborate set of oxidative stress conditions is displayed in Supplemental Table S8). Grey cells in the heatmap indicate missing values. Asterisk indicates ROS marker genes (Willems et al., 2016). B. Grey bars report the number of ChIP-confirmed target genes that are responsive (|log_2_ FC| > 1 and adj. *P* value < 0.01) to a number of oxidative stresses (Supplemental Table S8) as indicated in the X axis. The line indicates the cumulative fraction. C. Tissue-specific expression profiles of *ANAC102* and its ChIP-confirmed target genes obtained from the Genevestigator 8.0.2 Condition Search Tool (Hruz et al., 2008). Part of the heatmap focussing on rosette and root tissues is displayed.
